# Functions of the AP2/ERF family transcription factor AIL7 in immunity against soilborne clubroot pathogen in Arabidopsis

**DOI:** 10.1101/2024.05.22.595381

**Authors:** Kethmi Nirmani Jayawardhane, Tharangani Somarathna, Victor P. Manoli, Sheau-Fang Hwang, Stephen E. Strelkov, Stacy D. Singer, Guanqun Chen

**Author notes:** To whom correspondence should be addressed: Guanqun Chen [;] and Stacy D. Singer [;]. The author(s) responsible for distribution of materials integral to the findings presented in this article in accordance with the policy described in the Instructions for Authors (https://academic.oup.com/plcell/pages/General-Instructions) is: Guanqun Chen.

## Abstract

Soilborne pathogens can be highly devastating, and clubroot, caused by *Plasmodiophora brassicae*, is particularly destructive to cruciferous plants. Although many AP2/ERF family transcription factors (TFs) have crucial physiological functions, very little is known regarding their functions in the context of soilborne diseases. Here we investigated the roles of AINTEGUMENTA-LIKE 7 (AIL7), an AIL sub-family TF in the AP2/ERF family, in plant immunity against clubroot. Unexpectedly, both *AIL7* overexpression and mutant Arabidopsis lines exhibited increased tolerance to *P. brassicae*. Subsequent analysis revealed significant transcriptional alterations in genes linked to pathogen response, along with notable differences in genes associated with salicylic acid (SA) and jasmonic acid (JA) defense pathways, compared to wild-type plants. Interestingly, there was a tendency for up-regulation of SA- and JA-related genes in *AIL7* overexpression and mutant lines in the absence, rather than presence, of *P. brassicae*. Subsequent phytohormone analyses confirmed these results. Taken together, AIL7 has an important role in maintaining constitutive systemic acquired resistance, involving phytohormone mediated defense, and this, rather than an accumulation of SA following *P. brassicae* challenge, primes the plants for improved clubroot resistance, which would shed light on exploring the functions of other AP2/ERF family TFs in plant immunity against soilborne pathogens.

## Introduction

Soilborne plant pathogens can be highly devastating in terms of crop health, yield, and economic sustainability, and the chemical strategies currently deployed to combat these pathogens have often proven inadequate (Delgado-Baquerizo et al., 2020). Clubroot, for example, is one of the most destructive soilborne diseases of plants in the Brassicaceae family, which provides the second-largest source of vegetable oil and third largest source of vegetables, including rapeseed, mustard, radish, cabbage, kohlrabi, cauliflower, and broccoli (Gan et al., 2022; Hasan et al., 2021; FAO 2020). This disease is caused by an obligate biotrophic protist, *Plasmodiophora brassicae*, which can survive in the soil as long-lived resting spores. Clubroot is characterized by the formation of conspicuous galls or clubs on the roots of infected plants, resulting from cell hyperplasia and hypertrophy. These galls hamper the normal absorption of water and nutrients, resulting in stunted aboveground growth and reduced yields (Khalid et al., 2022; Dixon, 2009). The economic significance of Brassicaceae crops highlights the importance of understanding the dynamics of clubroot disease and its regulation at the molecular level. Furthermore, the ability of *P. brassicae* to infect a wide range of Brassicaceae hosts positions it as an interesting model for studying plant immunity to soilborne diseases.

Plants rely on complex defense mechanisms to recognize and defend themselves against potential stresses (Dodds et al., 2024). This inherent immunity can manifest as either pre-existing or induced defense strategies. Pre-existing defenses are general, innate responses that include chemical compounds with antimicrobial properties and physical barriers, while induced defenses arise from the detection of specific pathogen-associated or host-derived molecular patterns (microorganism-associated molecular patterns {MAMPs}, damage-associated molecular patterns {DAMPs}), or microbial effectors, which include both apoplastic and intracellular effectors (Dodds et al., 2024; Wang et al., 2022). The detection of MAMPs, DAMPs, or microbial effectors by plant cell surface receptors, known as pattern recognition receptors (PRRs), initiates pattern-triggered immunity (PTI) (Dodds et al., 2024), which broadly counters a wide range of microbes. When pathogens breach PTI, plants deploy effector-triggered immunity (ETI), providing a more specific response mediated by resistance (R) proteins, which are typically encoded by nucleotide-binding site and leucine-rich repeat (NBS-LRR or NLR) proteins that are activated through direct or indirect interactions with pathogen effectors (Dodds et al., 2024).

The recognition of a pathogen through specific interactions between elicitors (pathogen- or host-derived molecules) and host receptors initiates an intracellular signaling process that process activates downstream effector enzymes, generating specific secondary messengers that trigger various stress response signaling cascades (Dodds et al., 2024). Such cascades include the rapid build-up of reactive oxygen species (ROS), shifts in cellular ion flux, activation of mitogen-activated protein kinase (MAPK) cascades, changes in gene expression, and the production of stress-related hormones such as salicylic acid (SA), jasmonic acid (JA), and ethylene (Dodds et al., 2024; Zhou & Zhang, 2020). These signaling pathways lead to modifications in post-translational regulation and transcriptional reprogramming through the function of microRNAs (miRNAs), long non-coding RNAs (lncRNAs), and transcription factors (TFs) (Zhou & Zhang, 2020), which influences the expression and activity of numerous defense-related genes, including those encoding pathogenesis-related (PR) proteins (Dos Santos & Franco, 2023). While these defense responses are induced in a localized manner, they also trigger the spread of resistance throughout the plant in a process termed systemic acquired resistance (SAR) (Wang et al., 2022).

TFs play a critical role in orchestrating plant immune responses, acting as key regulators in the complex network of defense mechanisms (Feng et al., 2020; Ng et al., 2018). Among many diverse families of TFs, the APETALA2/ETHYLENE RESPONSIVE FACTOR (AP2/ERF) family stands out for the multifaceted roles its members play in plant development, physiology, and stress response (Feng et al., 2020). Although the majority of studies involving members of this family have focused on abiotic stresses such as cold and drought, recent studies have shown that AP2/ERFs also play an important function in resistance to phytopathogens and insect pests (Nie & Wang, 2023). Indeed, members of the AP2/ERF family recognize GCC boxes and activate or inhibit the expression of PR genes, and may also be involved in hormonal regulation related to defense response (Dong et al., 2015; Feng et al., 2020; He et al., 2001; Krizek et al., 2016; Licausi et al., 2013; Ma et al., 2024; Nie & Wang, 2023; Zhao et al., 2017). However, their functions and associated molecular mechanisms in root-related diseases are largely unknown. As such, exploring the functions of AP2/ERF TFs in terms of soilborne diseases holds promise for providing novel insight into plant immune mechanisms and could enhance our ability to develop targeted strategies for crop protection.

TFs in the AINTEGUMENTA (AIL) subfamily are important members of the AP2/ERF family, possessing two AP2 DNA binding domains approximately 60-70 amino acids in length (Horstman et al., 2016). In Arabidopsis, there are 8 *AIL* genes, including *AINTEGUMENTA* (*ANT*), *AIL1, BABY BOOM* (*BBM*)/ *AIL2* and *PLETHORA1* (*PLT1*)/ *AIL3, PLT2/ AIL4*, *PLT3/AIL6*, *PLT5/ AIL5,* and *PLT7/AIL7*. All of these genes are expressed in dividing tissues, where they act as master regulators of developmental processes including embryogenesis, flower development, stem cell niche specification, meristem maintenance, organ positioning, and growth (Aida et al., 2004; Horstman et al., 2016; Krizek, 2009, 2015; Pinon et al., 2013), possibly through the regulation of phytohormone-related pathways (Krizek et al. 2021; Santuari et al. 2016). Recent studies have also reported functions for *AIL* genes in biotic stress tolerance, as evidenced by the enhanced *Pseudomonas syringae* resistance observed in *Arabidopsis thaliana* (Arabidopsis) *antail6* double mutants (Krizek et al., 2016).

In addition, our recent transcriptomic analysis also suggests the potential involvement of AIL7 in biotic stress response (Singer et al., 2021). AIL7 is known to function alongside ANT and AIL6 in floral organ initiation and determination (Han & Krizek, 2016; Krizek et al., 2016, 2020; Mudunkothge & Krizek, 2012), as well as with AIL5 and AIL6 in the positioning and outgrowth of lateral shoot and root organs (Hofhuis et al., 2013; Prasad et al., 2011; Santuari et al., 2016). However, the seed-specific overexpression of *AIL7* in Arabidopsis also led to significant alternations in many defense-related genes, including the upregulation of a large number of NBS-LRR and PR genes in developing siliques without pathogen inoculation (Singer et al., 2021). While PR genes typically encode enzymes that play a crucial role in the first line of defense against plant pathogens (Dos Santos & Franco, 2023), NBS-LRR genes encode a large class of R proteins that are key components of ETI (Kopec et al., 2021; Yu et al., 2017). Furthermore, these lines also exhibited the upregulation of numerous genes involved in SA and JA-related pathways in developing siliques (Singer et al., 2021). Collectively, these results hint at the possibility that *AIL7* may function in Arabidopsis immune response; however, whether AIL7, or more broadly AP2/ERF family TFs, function in the response against soilborne root pathogens is not known.

Based on these previous studies, we hypothesized that AIL7 may play a role in the molecular regulation of plant immunity against the clubroot pathogen. To test this hypothesis, we assessed the performance of *AIL7* constitutive overexpression and T-DNA mutant Arabidopsis lines following inoculation with *P. brassicae*. Additionally, we conducted further in-depth assessments of these plants at the metabolic and molecular levels to elucidate the precise regulatory mechanisms driving improved clubroot resistance in these lines. The results of this study expand our knowledge of the functions of AIL7 in clubroot resistance, and provide an additional genetic resource for the development of Brassicaceae crops with improved resistance downstream. Additionally, since AIL7 belongs to the AP2/ERF family of TFs, findings from this study also highlight the importance of further assessing other TFs in this family for their roles in plant immunity against soilborne root diseases.

## Results

### Characterization of Arabidopsis AIL7 constitutive overexpression and mutant lines

To analyze the functions of *AIL7* in clubroot disease resistance, we generated single-copy transgenic homozygous Arabidopsis lines constitutively overexpressing *AIL7*, and characterized *ail7* T-DNA Arabidopsis mutants (**Figure 1**). The three selected homozygous overexpression lines (OE2, OE5, OE12) exhibited 12.5-, 10.5-, and 8.2-fold increases in *AIL7* expression in roots compared to wild-type controls, respectively (**Figure 1A**). The two mutants, *ail7-1* (SAIL_1167_C10; termed M1 in this study) and SAIL_267_G12 (termed M2 in this study), were confirmed to be homozygous for T-DNA insertion sites within the 6^th^ exon and 5’UTR region, respectively (**Figure 1B, 1C**). The *ail7-1* mutant (M1) bears an interrupted coding sequence, which likely leads to the production of a partial and non-functional protein and is considered to be a knockout (Krizek, 2009). SAIL 267 G12 (M2), which is identified as a knockout T-DNA mutant for *AIL7* in the Arabidopsis Biological Resource Center, has not been studied previously. Homozygous lines of both mutants were used in subsequent experiments.

**Figure 1.**
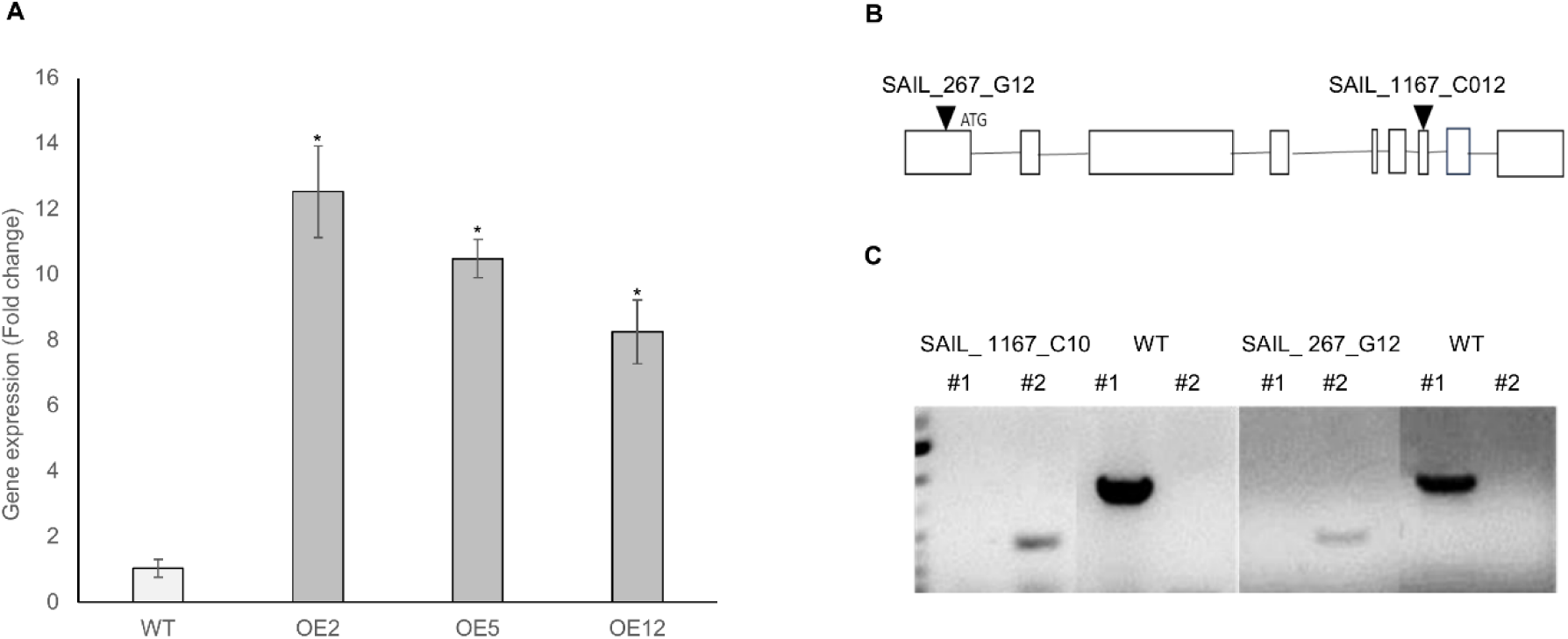
*AIL7* overexpression and T-DNA mutant line identification and characterization. (A) Relative expression of Arabidopsis *AIL7* in *AIL7* overexpression and wild-type (control) lines. Dark-colored blocks represent overexpression lines while light-colored blocks represent wild-type plants. Each block represents 2 biological replicates, and each replicate was prepared from homogenized root samples from approximatley 15 lines. (B) Schematic diagrams of the positions of T-DNA insertions in *ail7* single mutant alleles. Start and stop codons are indicated. (C) Identification of T-DNA insertion lines by PCR. LP and RP primers are located at the left and right sides of the T-DNA insertion, respectively. LBb1.3 was used as the flanking primer. In each gel image, #1 is the PCR sample with primers LP+RP and #2 is the PCR sample with primers LB1.3 +RP. OE, Arabidopsis *AtAIL7* constitutive overexpression lines; WT, wild-type Arabidopsis lines.

### AIL7 overexpression lines exhibit increased plant size, vegetative biomass, seed size and weight

To determine whether *AIL7* overexpression or mutation impacts plant growth and yield characteristics under non-stressed conditions, we carried out detailed analyses of *AIL7* overexpression and knockout mutant lines relative to wild-type Columbia 0 (Col-0) plants. In the case of *AIL7* overexpression lines, evaluation of seedling growth on ½ strength MS medium indicated that no significant changes in root or hypocotyl growth were present 10 days post-germination in any line except for AIL7-OE2 (**Figures 2A, 2B, 2C,** and **2D**). While there was a slight, but significant increase in root growth of AIL7-OE2 lines compared to wild-type **(Figures 2C and 2D)**, the fact that the majority of overexpression lines did not exhibit this same characteristic suggests that *AIL7* overexpression or mutation does not substantially alter seedling growth. Conversely, 25 days post-germination on potting mix, *AIL7* overexpression lines were noticeably larger than wild-type plants **(Figure 3A)**. Indeed, rosette diameter **(Figure 3B)** and leaf biomass per plant **(Figure 3C)** of *AIL7* overexpression lines were significantly greater than wild-type plants. Furthermore, the height of *AIL7* overexpression lines was significantly higher than wild-type plants, with lines OE2, OE5, and OE12 showing relative increases of 27.5%, 22.3% and 35.5%, respectively, compared to wild-type plants **(Figures 3D and 3E)**. Although we did not observe any significant alterations in the general appearance of *AIL7* overexpression seeds **(Figure 4A),** we did note a significant increase in seed area **(Figure 4B)** and 100 seed weight in the *AIL7* overexpression lines compared to wild-type **(Figure 4C)**. However, seed yield per plant was unchanged in *AIL7* overexpression lines compared to wild-type plants **(Figure 4D)**. In contrast, no significant alterations were observed in any of the above-mentioned characteristics in the *ail7* mutants compared to the wild-type **(Figures 2, 3 and 4)**.

**Figure 2.**
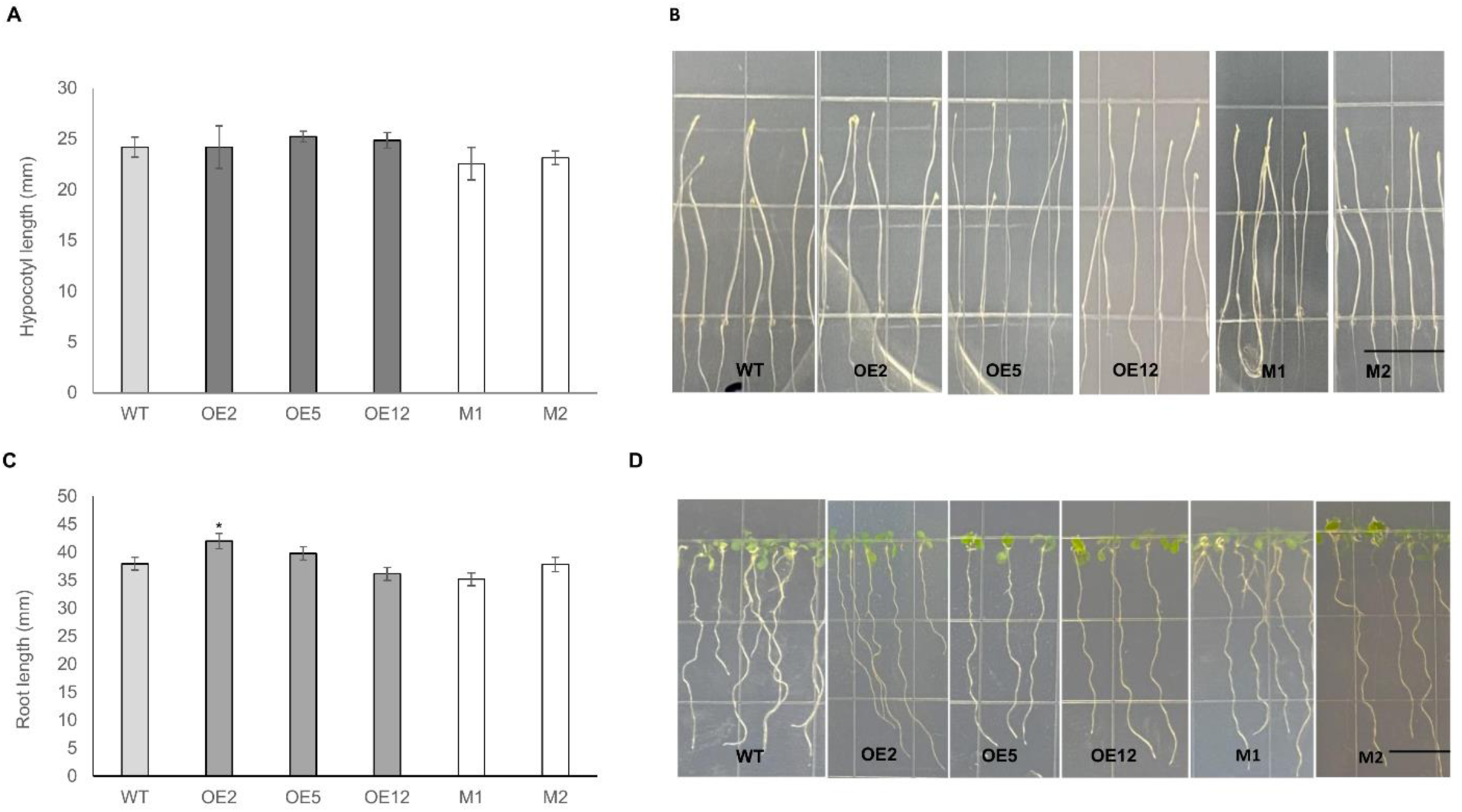
Morphology of *AIL7* overexpression, mutant and wild-type seedlings 10 days post germination. (A) The average length of dark-grown *AIL7* overexpression, mutant and wild-type hypocotyls where each bar represents approximately 20 hypocotyls and the picture shown in (B) is representative of three experiments conducted with similar results. (C) The average root length of *AIL7* overexpression, mutant and wild-type lines where each bar represents approximately 20 seedlings and the picture shown in (D) is representative of three experiments conducted with similar results. The values (A and C) are means ± SE of approximately 20 seedlings and the asterisks indicate significant differences at *P* < 0.05 (*) compared to wild-type, based on two-tailed student t-tests. Scale bars represent 1 cm. OE, Arabidopsis *AIL7* constitutive overexpression lines; M, *AIL7* mutant; WT, wild-type Arabidopsis lines.

**Figure 3.**
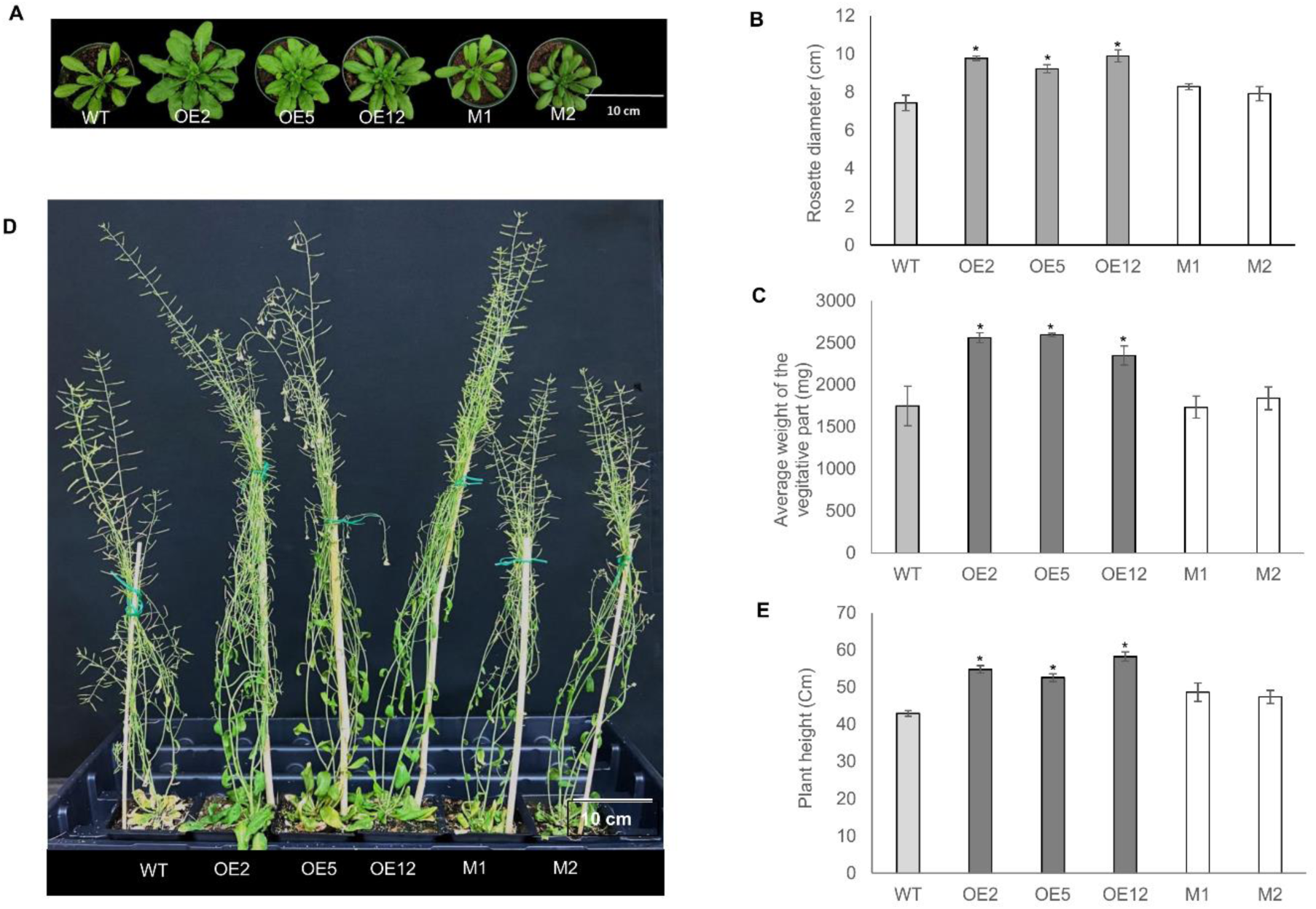
Phenotypes of *AIL7* overexpression, mutant and wild-type control plants grown in soil. (A) Phenotypes of Arabidopsis rosettes 25 days after sowing (B) Rosette diameter and (C) vegetative biomass. (D) Phenotype and (E) height of Arabidopsis plants 28 days after bolting. Each bar represents measurements of 10-12 induvial homozygous plants. Significant differences compared to the control are indicated by asterisks (*P*< 0.05). OE, Arabidopsis *AIL7* constitutive overexpression lines; M, *AIL7* mutants; WT, wild-type Arabidopsis lines.

**Figure 4.**
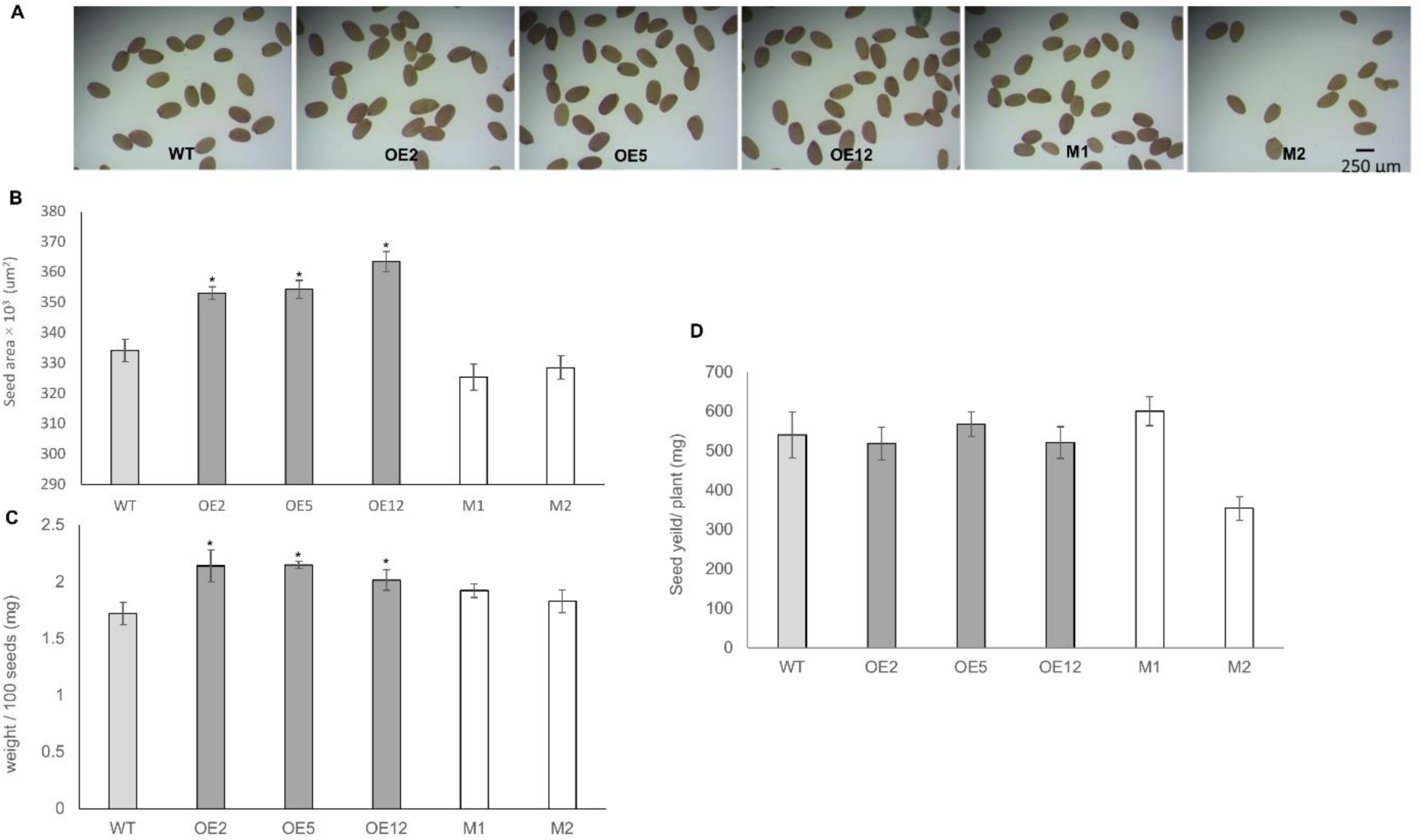
Morphology of *AIL7* overexpression, mutant and wild-type seeds. (A) Representative image of *AIL7* overexpression, mutant and wild-type seeds where (B) the seed size, (C) weight per 100 seeds, and (D) seed weight per plant were measured using 10 induvial plant lines for each homozygous overexpression, mutant and wild-type line. Significant differences compared to the control is indicated by an asterisk (*P* < 0.05). Scale bars represent 200 µm. OE, Arabidopsis *AIL7*constitutive overexpression lines; M, *AIL7* mutant WT, wild-type Arabidopsis lines.

### AIL7 plays a role in the response of Arabidopsis to P. brassicae

To examine the role of *AIL7* in clubroot disease response, we first evaluated disease severity in homozygous *AIL7* overexpression lines, T-DNA insertion *ail7* knockout mutants, and wild-type Arabidopsis, following inoculation with *P. brassicae* pathotype 3H at a concentration of 1.0 x 10^5^ resting spores/mL water. Plants began to show a purpling of the leaves approximately 16-18 days post-inoculation **(Figure 5A and 5B),** and root disease severity measurements were carried out 21 days post-inoculation (**Figure S1A**; Zhou et al., 2022). Both the overexpression and knockout mutants had significantly lower disease indexes (DI) (41.3% - 62.1% and 39.9 - 66.7%, respectively) than wild-type plants (72.71%), indicating enhanced resistance to *P. brassicae* (**Figure 5C**). We also performed disease severity assessments following inoculation with a higher inoculum concentration (1.0 x 10^6^ resting spores/mL water). While a similar trend was observed in terms of DI values, the differences were not significant **(Figure S1B)**, which led us to work with the lower spore concentration as reported in other studies (Zahr et al., 2022). Collectively, these results indicate that *AIL7* plays a role in clubroot resistance in Arabidopsis.

**Figure 5.**
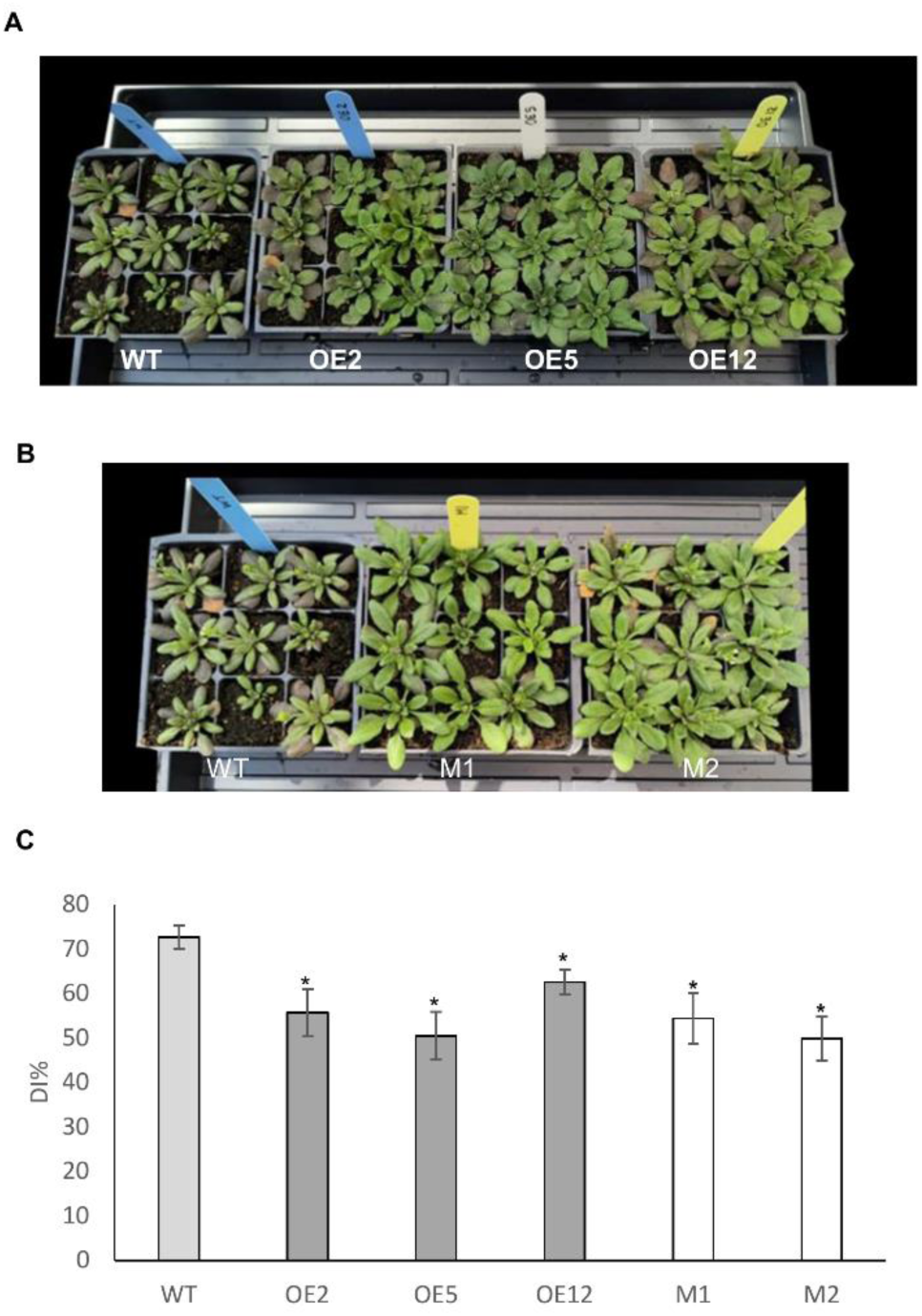
Evaluation of *AIL7* overexpression, mutant and wild-type Arabidopsis plants in response to inoculation with *P. brassicae* pathotype 3H. (A) Above-ground phenotypic observations of inoculated *AIL7* overexpression (B) *AIL7* mutant plants compare to wild-type control plants16 days after inoculation (C) Disease index of *AIL7* overexpression, mutant and wild type Arabidopsis lines 21 days after inoculation with *P. brassicae* inoculum (1*10^5^ spores/ mL). The individual severity ratings were used to calculate a disease index in each case. Values are means ± SE of four independent replicates with 20-25 plants per replicate. Asterisks indicate significant differences at *P* < 0.05 (*) compared to wild-type plants, based on two-tailed student t-tests. OE, Arabidopsis *AIL7* constitutive overexpression lines; M, *AIL7* mutant WT, wild type Arabidopsis lines.

### AIL7 overexpression and mutant lines display alterations in SA and JA levels compared to wild-type plants

Since the involvement of *AIL* genes in growth regulatory processes and stress tolerance in many cases has been shown to be related to phytohormones (Bauer et al., 2021; Ding et al., 2018; Krizek et al., 2020; Santuari et al., 2016), we hypothesized that the effect of *AIL7* on clubroot resistance is also linked to phytohormone biosynthesis and/or modification. Thus, we measured SA, JA, and auxin levels in the roots of *AIL7* overexpression, mutant, and wild-type Arabidopsis lines following *P. brassicae-* or mock-inoculation, to elucidate how *AIL7* influences phytohormone levels (**Figure 6**). In the absence of *P. brassicae*, both *AIL7* overexpression and mutant lines demonstrated significantly elevated SA content (0.40 ± 0.08 nmol/g and 0.04 ± 0.06 nmol/g respectively) compared with the wild-type (0.16 ± 0.03 nmol/g). In contrast, after inoculation with *P. brassicae*, no significant differences in SA levels were observed in either *AIL7* overexpression (1.65 ± 0.4 nmol/g) or mutant lines (1.27 ± 0.09 nmol/g) relative to the wild-type (1.21 ± 0.2 nmol/g) **(Figure 6A)**. These findings indicate that *AIL7* may play a role in the modulation of SA metabolism in Arabidopsis, and that it is the increased SA levels observed in *AIL7* overexpressing and mutant lines in the absence of the pathogen, rather than SA buildup following infection, that might contribute to clubroot tolerance.

**Figure 6.**
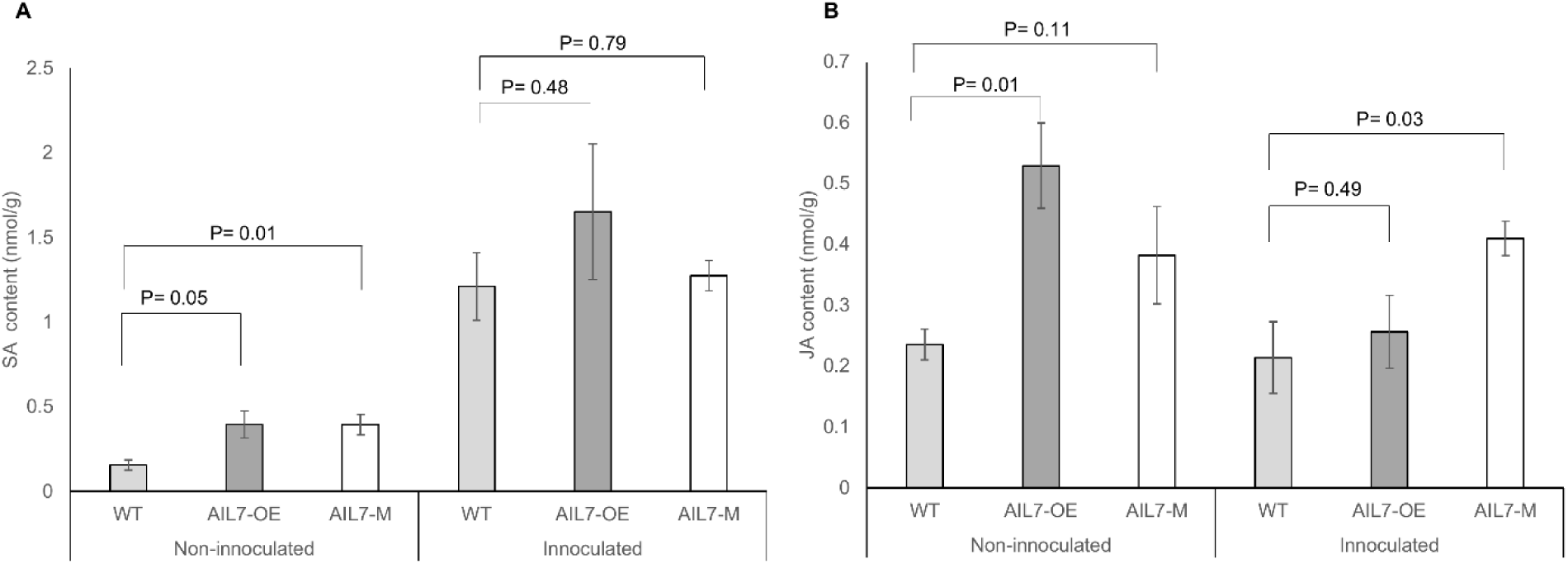
SA (A) and JA (B) content in *P. brassicae* inoculated and non-inoculated Arabidopsis roots. Error bars denote ± SE of four independent replicates with approximately 20 plants for each replicate. P values are indicated in each comparison and *P* ≤ 0.05 are considered significant differences compared to wild-type, based on 2-tailed student t-tests. AIL7-OE, Arabidopsis *AIL7* constitutive overexpression lines; AIL7-M, *AIL7* mutant WT, wild type Arabidopsis lines.

Interestingly, in the absence of *P. brassicae* inoculation, JA content was also significantly higher in *AIL7* overexpression lines (0.53 ± 0.07 nmol/g) compared to wild-type plants (0.24 ± 0.02 nmol/g). Although there was a similar trend in *ail7* mutants (0.38 ± 0.08 nmol/g), these differences were not significant (**Figure 6B**). Conversely, following inoculation with *P. brassicae*, JA levels were significantly greater in *ail7* mutant lines (0.41 nmol/g), but not in *AIL7* overexpression lines, relative to the wild-type (0.2 nmol/g; **Figure 6B**). These results suggest that enhanced levels of JA in *AIL7*-modulated plants may also contribute to clubroot tolerance.

Since previous studies have linked members of the AIL family to auxin biosynthesis (Krizek et al., 2016; Pinon et al., 2013), and auxin is believed to be involved in root gall development (Ludwig-Müller et al., 1996), we also examined auxin levels in the roots of *AIL7* overexpression and mutant lines, as well as the wild-type control, under inoculated and mock-inoculated conditions. However, unlike SA and JA, no significant differences in auxin levels were observed in *AIL7* overexpression or mutant lines compared to wild-type plants following either *P. brassicae-* or mock-inoculation (**Figure S2**).

### The overexpression and mutation of AIL7 in Arabidopsis leads to the transcriptional modulation of genes involved in disease resistance and hormonal regulation

To investigate how *AIL7* contributes to clubroot resistance at the transcriptional level, we analyzed the expression profiles of various immune response-related genes in root samples. Given that seed-specific overexpression of *AIL7* in Arabidopsis was previously reported to result in the up-regulation of various genes encoding putative R and PR proteins in developing siliques (Singer et al., 2021; Table S2), we began by assessing the expression of TIR-NBS-LRR genes (AT1G63860, AT5G45240, AT5G38340, AT4G36140, AT1G72950), PR thaumatin superfamily genes (AT1G18250, AT1G73620), and PR genes (*PR1, PR2, PR5*) (**Table 1**; **Table 2**). All of these genes have been linked to the clubroot immune response in previous studies (Adhikary et al., 2022; Fu et al., 2019; Irani et al., 2018; Jia et al., 2017; Singer et al., 2021; Zhou et al., 2022). While most of these genes did not show any significant changes in expression levels in the root tissues of *AIL7* overexpression lines following either *P. brassicae* or mock-inoculation (**Figures 7A and 7C**), both *PR1* and *PR2* were significantly upregulated in the T-DNA mutant lines compared to wild-type plants following mock-inoculation (**Figure 7A**). Conversely, *PR2* was downregulated in *AIL7* overexpression plants following *P. brassicae* inoculation **(Figure 7C)**. Expression of the TIR-NBS-LRR gene AT4G36140 was significantly down-regulated in both the overexpression and mutant lines compared to wild-type plants following *P. brassicae* inoculation (**Figure 7C**). Expression of AT1G63860 was not in the detectable range in either inoculated or mock-inoculated plants.

**Figure 7.**
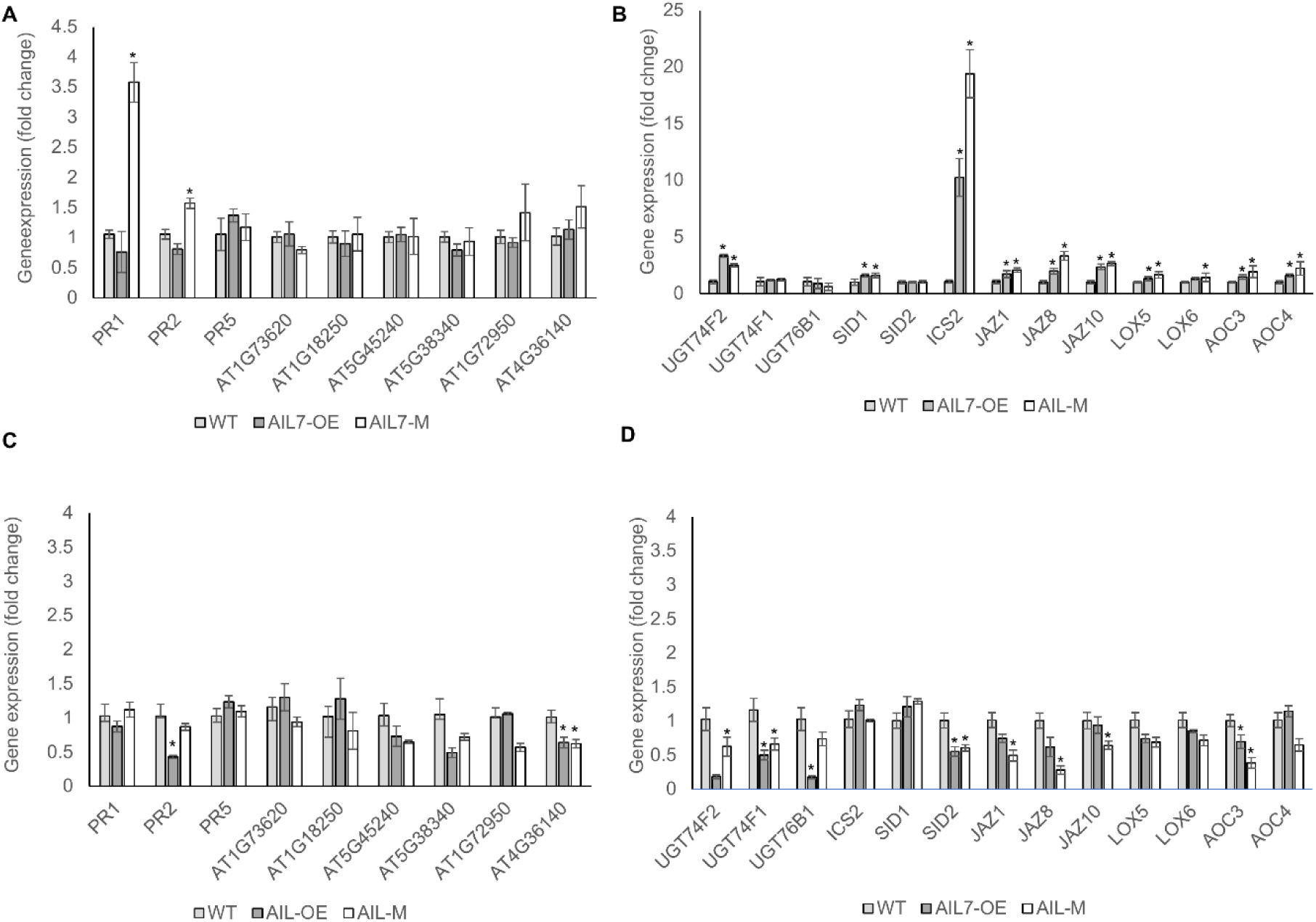
Expression levels of genes related to pathogenesis and hormone biosynthesis in root samples of Arabidopis lines inoculated with *P. brassicae*, along with mock-treated controls. (A) Defense-related and (B) hormone-related gene expression in non-inoculated plants and (C) defense-related (D) hormone-related gene expression in non-inoculated plants. Error bars represent ±SE of 4 biological replicates, each consisting of approximately 20 individual plants. Asterisks indicate a significant difference at *P* < 0.05 (*) compared to wild-type, based on 2-tailed student t-tests. AIL7-OE, Arabidopsis *AIL7* constitutive overexpression lines; AIL-M, *AIL7* mutant WT, wild-type Arabidopsis lines.

**Table 1.** Differentially expressed disease resistance genes in previous study of *AIL7* seed-specific overexpressed Arabidopsis plants (more than 10-fold increase)

| Gene ID | Ratio | p-value | Description |
| --- | --- | --- | --- |
| AT3G25510 | 10.5 | 0.000594 | Disease resistance protein (TIR-NBS-LRR class), putative |
| AT4G11170 | 11.33 | 0.027383 | disease resistance protein (TIR-NBS-LRR class), putative |
| AT2G15080 | 12.6 | 8.94E-09 | Disease resistance family protein |
| AT3G04220 | 12.67 | 0.000423 | Disease resistance protein (TIR-NBS-LRR class), putative |
| AT3G23110 | 15.33 | 0.026514 | Disease resistance family protein |
| AT5G38340 | 16.11 | 2.46E-18 | Disease resistance protein (TIR-NBS-LRR class), putative |
| AT1G63860 | 16.77 | 1.06E-15 | Disease resistance protein (TIR-NBS-LRR class) |
| AT1G57630 | 23.89 | 0.008219 | Disease resistance protein (TIR class), putative |
| AT5G45240 | 25.33 | 8.93E-11 | Disease resistance protein (TIR-NBS-LRR class), putative |
| AT3G24900 | 28.4 | 0.047233 | Disease resistance family protein / LRR family protein |
| AT3G25010 | 87.92 | 0.00245 | Disease resistance family protein |
| AT2G32680 | 224 | 3.84E-05 | Disease resistance family protein |
| AT1G18250 | 98.22 | 3.41E-11 | Pathogenesis-related thaumatin superfamily protein |
| AT1G73620 | 119.67 | 1.77E-15 | Pathogenesis-related thaumatin superfamily protein |
| AT1G75040 | 320.25 | 4.63E-25 | Pathogenesis-related gene 5 |
| AT2G14610 | 35.74 | 1.48E-21 | Pathogenesis-related gene 1 |
| AT3G57260 | 142.57 | 1.11E-05 | BGL2 (PATHOGENESIS-RELATED PROTEIN 2) |

**Table 2.**
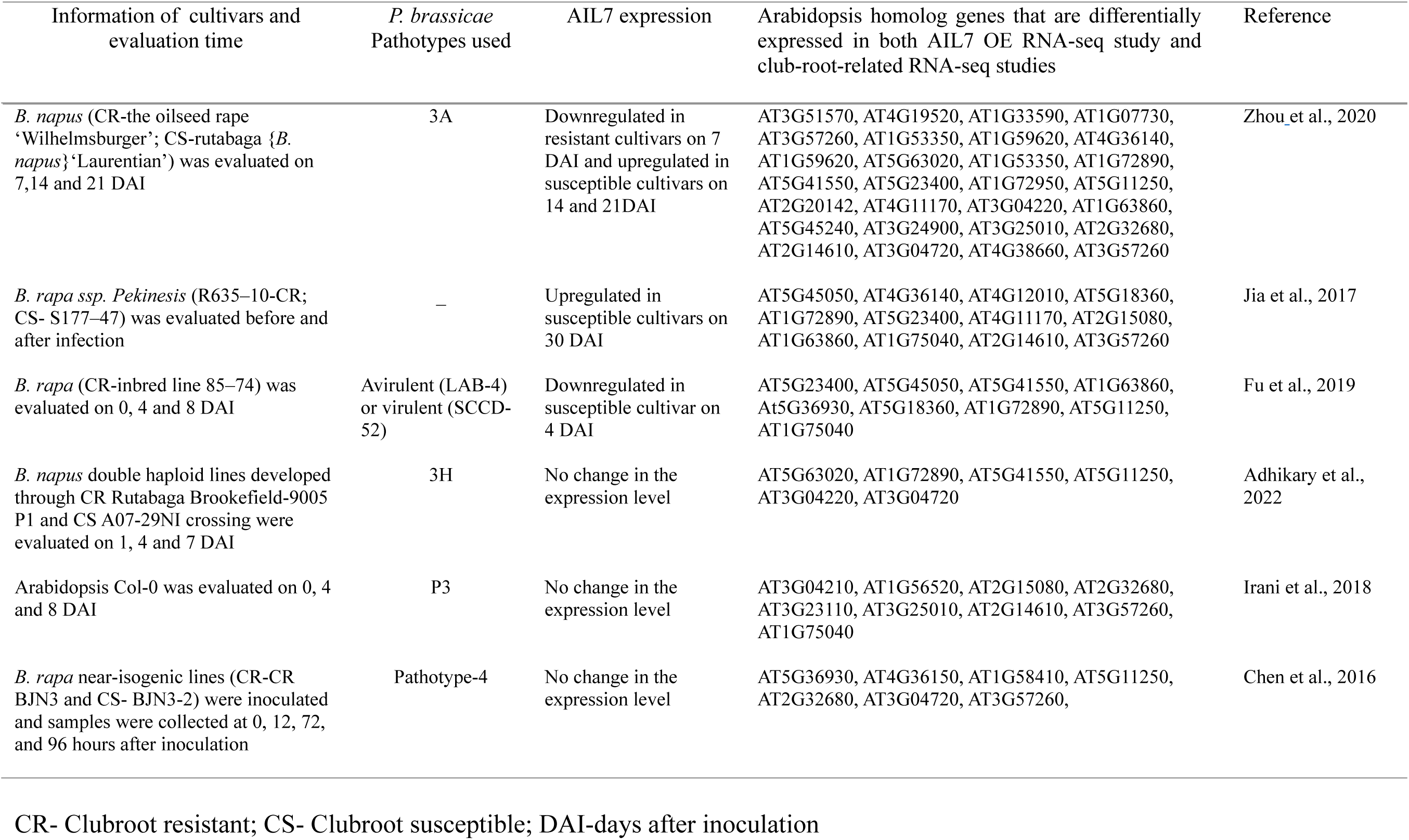
Previous RNA-Seq studies of clubroot resistance in Arabidopsis and Brassica *spp*.

Since we observed significant alterations in phytohormone levels in *AIL7*-modulated lines, we subsequently analyzed the expression of genes with known functions in phytohormone signaling in *AIL7* overexpression and mutant lines compared to the wild-type following *P. brassicae* inoculation or mock treatment (**Figures 7B an 7D**). In terms of SA biosynthesis in the absence of *P. brassicae* inoculum, the overexpression or knockout of *AIL7* did not lead to significant changes in the expression of *SALICYLIC ACID INDUCTION DEFICIENT 2* (*SID2*), which catalyzes the first step of SA biosynthesis to convert chorismate to isochorismate (Bauer et al., 2021; Ding et al., 2018). However, there was a significant increase in the expression of both *ISOCHORISMATE SYNTHESIS* (*ICS2*), which functions redundantly with *SID2* in chloroplasts, and *SID1*, which is involved in the export of isochorismate to the cytosol, compared to wild-type (**Figure 7B**), suggesting that SA biosynthesis may be constitutively activated in the absence of disease pressure in these lines. Similarly, we observed the significant upregulation of *UGT74F2* in both mutant and overexpression lines. This gene serves an important function in the glucosylation of SA to SA-glucose ester (Peng et al., 2021), and its upregulation suggests that in the absence of *P. brassicae*, SA may undergo a higher extent of modification to its storage form (**Figure 7B**). However, no significant differences were noted in the expression of *UGT74F1* or *UGT76B1*, which are also involved in SA glucosylation (Peng et al., 2021), following mock treatment (**Figure 7B**).

Following inoculation with *P. brassicae*, on the other hand, the expression of genes related to SA biosynthesis and modification exhibited distinct expression profiles (**Figure 7D**). In this case, *SID2* expression levels were significantly reduced in both *AIL7* overexpression and mutant lines compared to wild-type, while no significant differences were observed in *SID1* or *ICS2* expression **(Figure 7D)**. Similarly, *UGT74F1* was significantly down-regulated in both *AIL7* overexpression and mutant lines compared to wild-type plants, and while the same trend was noted for *UGT74F2* and *UGT76B1*, their downregulation was significant only in *AIL7* overexpression lines **(Figure 7D)**. Taken together, the expression analysis of SA-related genes was consistent with our phytohormone data, where we observed constitutively high amounts of SA in *AIL7* overexpression and mutant plants compared to wild-type.

In terms of JA biosynthesis and signaling in the absence of *P. brassicae* inoculation, several genes also exhibited differential expression levels in *AIL7* overexpression and mutant lines compared to wild-type **(Figure 7B)**. Two LOX family genes (*LOX5* and *6*), which catalyze the first step in JA biosynthesis (converting linolenic acid to 13-hydroperoxylinolenic acid) (Ruan et al., 2019), were significantly upregulated in *AIL7-*overexpressing plants compared to wild-type **(Figure 7B)**. The same trend was noted in *ail7* mutant plants; however, the difference was significant only in the case of *LOX5* (**Figure 7B**). Moreover, *ALANINE OXIDASE CYCLASE* family genes (*AOC3* and *4*), which catalyze the essential cyclization of alanine oxide to form *cis*- (+)-OPDA during JA biosynthesis (Ruan et al., 2019) were also significantly upregulated in both *AIL7* overexpression and mutant lines compared to wild-type (**Figure 7B**). These results suggest that in addition to SA biosynthesis and modification, JA biosynthesis is also affected in *AIL7*-modified lines. The JASMONATE-ZIM DOMAIN protein-encoding genes *JAZ1*, *8,* and *10*, which function as repressors of JA signaling (Liu & Timko, 2021), were all also upregulated in both *AIL7* overexpression and mutant lines following mock treatment.

In contrast, following inoculation with *P. brassicae*, there was a trend for the downregulation of JA-related gene expression in *AIL7* overexpression and mutant lines compared to wild-type plants (**Figure 7D**). These differences, however, were not always significant. Indeed, in terms of JA biosynthetic genes, only *AOC3* transcript levels were significantly reduced in both the overexpression and mutant lines compared to wild-type plants **(Figure 7D)**, while the expression of all three *JAZ* genes assessed (*JAZ1*, *JAZ 8* and *10*) was significantly downregulated in *ail7* mutant, but not overexpression, lines **(Figure 7D)**. Collectively, these findings suggest that the regulation of JA signaling in *AIL7* overexpression and mutant plants is likely complex, as there are simultaneous effects on the transcription of JA biosynthetic and JA-response repressor genes.

### The overexpression of AIL7 in Arabidopsis reduces the transcript levels of other AIL family members

Previous studies have indicated that AIL7 shares a similar DNA-binding motif with three other AIL family members, including AIL3, AIL4 and AIL6 (Malley et al., 2016; Santuari et al., 2016; Singer et al., 2021). As such, we hypothesized that the similarity in clubroot response observed between *AIL7* overexpression and mutant lines may reflect an unexpected transcriptional alteration in one of these *AIL* genes. To elucidate any such effect, we examined the expression profiles of *AIL3*, *AIL4*, *AIL5* and *AIL6* in the roots and leaves of *AIL7* overexpression and mutant lines compared to wild-type plants. No significant differences in expression were observed for any of the tested genes in the roots of *AIL7* overexpression or mutant lines compared to wild-type (**Figure 8A**), suggesting that the modulation of *AIL7* did not affect the expression of these genes in roots. However, *AIL7* overexpression led to a reduction in the transcript levels of *AIL6* and *AIL5* in leaves (**Figure 8B**).

**Figure 8.**
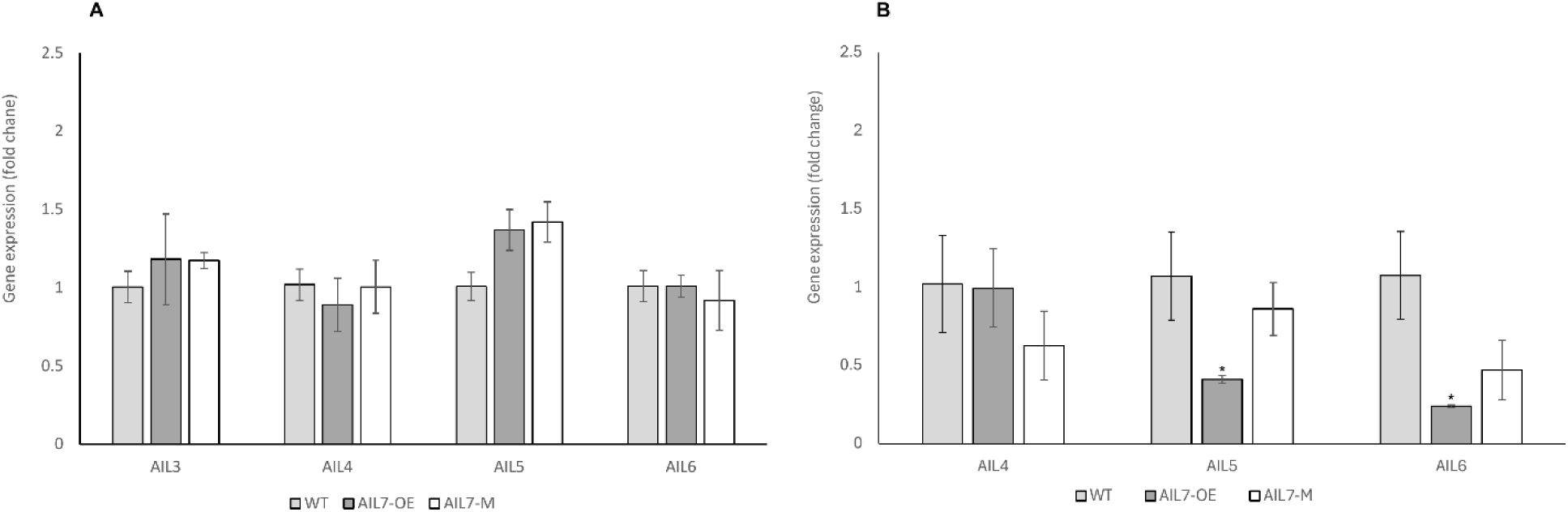
Expression levels of *AIL* family members in *AIL7* overexpression, mutant and wild-type roots (A), and leaves (B). Error bars represent ±SE of 3-4 biological replicates. AIL7-OE, Arabidopsis *AIL7* constitutive overexpression lines; AIL-M, *AIL7* mutant WT, wild-type Arabidopsis lines. The asterisks indicate a significant difference at *p* ≤ 0.05 (*) and compared with the wild-type, based on the 2-tailed student t-test.

## Discussion

Genes belonging to the *AIL* family are predominantly active in growth-centric regions of roots and shoots, crucially influencing the formation and layout of organ primordia (Hofhuis et al., 2013; Pinon et al., 2013), and there is a growing body of evidence suggesting that these genes might also participate in defense-related pathways (Krizek et al., 2016; Santuari, et al., 2016; Singer et al., 2021). In a previous study, the seed-specific overexpression of *AIL7* in Arabidopsis led to significant alterations in the expression of many defense-related genes in developing siliques in the absence of pathogen pressure (**Table 1**; **Table S2**; Singer et al., 2021). Since the role of *AIL* genes in plant defense against soilborne pathogens, including clubroot, has not yet been investigated, we initially sought to explore this transcriptomic data further in the context of clubroot disease.

Interestingly, many of the disease resistance-related, differentially expressed genes (DEGs) identified in these seed-specific *AIL7* overexpression Arabidopsis lines in the absence of *P. brassicae*-inoculation (**Table 1**) overlapped with DEGs observed previously in *Brassicaceae* species inoculated with *P. brassicae* compared to mock-treated controls, including those encoding several TIR-NBS-LRR and PR proteins (**Table 2**; Adhikary et al., 2022; Chen et al., 2016; Fu et al., 2019; Irani et al., 2018; Jia et al., 2017; Zhou et al., 2020). For instance, homologs of more than 20 transcripts with significantly altered levels in the developing siliques of *AIL7* seed-specific overexpression Arabidopsis lines were also categorized as DEGs in a transcriptomic comparison of clubroot susceptible/resistant *Brassica napus* cultivars at 3 different time-points in *P. brassicae*-inoculated plants compared to mock-inoculated plants (**Table 2**; Zhou et al., 2020). Furthermore, the analysis of multiple transcriptomic datasets comparing clubroot-resistant and susceptible *Brassica* spp. infected with *P. brassicae* revealed that *AIL7* expression levels were substantially different in resistant/susceptible cultivars compared to non-inoculated plants; however, the patterns observed were not consistent among studies (**Table 2**). For example, *AIL7* was found to be up-regulated in clubroot-susceptible *B. napus* and *B. rapa* cultivars following *P. brassicae* inoculation in several studies (Zhou et al., 2020; Jia et al., 2017). In contrast, *B. rapa* has been found to exhibit a reduction in *AIL7* expression 4 days post-inoculation in clubroot-susceptible cultivars (Fu et al., 2019), while no significant changes in *AIL7* expression were noted between susceptible and resistant *B. napus* or *B. rapa* lines following *P. brassicae* inoculation in other studies (Adhikary et al., 2022; Chen et al., 2016). Taken together, these previous results indicate that while *AIL7* may play a potential role in clubroot resistance, further research will be required to unravel its precise transcriptional effects under different growth and inoculation regimes. In the current study, we evaluated the clubroot susceptibility of Arabdidopsis *AIL7* constitutive overexpression and knockout lines to explore the function of *AIL7* in clubroot response and expand our understanding of the AP2/ERF family TFs in the context of soilborne pathogens in general.

In the present study, *AIL7* overexpression plants displayed enhanced plant, seed size and weight compared to wild-type plants (**Figures 3 and 4**). It is well-known that AIL TFs are involved in plant growth and development, and several studies have implied their involvement in organ size determination in different plant species (Ding et al., 2018; Krizek, 2009; Nole-Wilson et al., 2005; Wang et al., 2022). For example, the ectopic expression of *BrANT-1* in Arabidopsis caused an enhancement in organ size and affected plant height, as well as leaf, silique, and seed size, compared to wild-type plants by promoting cell proliferation (Ding et al., 2018). Similarly, the ectopic expression of *AIL5* in Arabidopsis resulted in increased floral organ size (Nole-Wilson et al., 2005), whereas the overexpression of *AIL2* in *B. napus* also promoted cell proliferation (Boutilier et al., 2002). Thus, it is likely that *AIL7* may also have a similar effect in this context. The fact that the *ail7* mutants did not exhibit any abnormal phenotype in our study (**Figures 2, 3 and 4**), which corresponds with previous findings with this gene (Nole-Wilson et al., 2005; Krizek, 2009), can almost certainly be attributed to genetic redundancy of *AIL* family members. This aligns with the fact that at least certain AIL family members can bind the same DNA motifs and regulate overlapping gene sets, leading to partial compensation when certain family members are absent (Santuari et al., 2016).

Intriguingly, both *AIL7* overexpression and mutant lines exhibited a significant reduction in DI compared to wild-type plants following inoculation with *P. brassicae* pathotype 3H (**Figure 5C**), which indicates enhanced disease resistance. Genetic redundancy within a gene family can in certain instances allow for compensation among mutants (Peng, 2019), where the overexpression of one gene can lead to the suppression of other family members, or vice versa (Tayengwa et al., 2020), resulting in analogous phenotypes in both knockout and overexpression scenarios (Prelich, 2012; Cusack et al., 2021). Indeed, previous research has demonstrated cross-transcriptional regulation among *AIL* family members in single and double mutants, which can affect overall phenotypic outcomes (Krizek et al., 2016) and suggests that complex interactions and regulatory processes within the *AIL* family could be contributing to similar disease resistance phenotypes in *AIL7* overexpression and mutant lines. While alterations in the expression of *AIL1*, *ANT*, *AIL2*, *AIL4*, *AIL5*, and *AIL6* were observed in the siliques of Arabidopsis seed-specific *AIL7* overexpression lines previously (Singer et al., 2021), we did not observe significant alterations in the transcript levels of *AIL3*, *AIL4*, *AIL5*, or *AIL6* in the roots of constitutive *AIL7* overexpression lines in the present study (**Figure 8A**). However, we did note a reduction in the expression of *AIL6* and *AIL5* in the leaves of our *AIL7* overexpression lines (**Figure 8B**). These findings suggest that the transcriptional cross-regulation of *AIL* genes is tissue-specific, and its effect in *AIL7* overexpression and mutant lines remains to be clarified.

It has also been shown previously that mutant and overexpression lines can exhibit similar phenotypes in plants when the overexpressed/mutated gene encodes a component of a multiprotein complex, possibly due to interference with the function of the respective complex (Prelich, 2012). In such protein-complex regulatory networks, the overexpression of a gene could activate a negative feedback loop, which suppresses the pathway downstream of the target gene and results in a phenotype resembling that of a loss-of-function mutant (Bao et al., 2022). For example, similar morphological deformities in pollen grains and reduced pollen viability were observed previously in tomato *ABORTED MICROSPORES* (*SlAMS*) mutants and overexpressing lines, which was suggested to occur due to the possible competitive transcriptional repression of upstream transcription factor-encoding genes involved in pollen development in overexpressing lines, thus leading to the disruption of the associated regulatory network (Bao et al., 2022). Since AIL7 is likely to function within a protein complex with feedback regulatory mechanisms (Horstman et al., 2016; Krizek et al., 2023) it is possible that similar mechanism is at play in our *AIL7* overexpression lines in the context of pathogen defense. A similar phenomenon has been observed with *POWDERY MILDEW RESISTANCE 4* (*PMR4*), which encodes a callose synthase, whereby both overexpressing and knockdown/knockout mutants have been demonstrated to display enhanced disease resistance against the causal pathogens of powdery mildew and downy mildew, as well as *Phytopthora infestans,* in several plant species. However, in this instance, such an effect was suggested to occur due to the dual function of this gene in callose deposition (Blümke et al., 2013; Nishimura, et al., 2003; Santillán Martínez et al., 2020; Vogel & Somerville, 1999). At present, it is unknown whether AIL7 possesses multiple, contrasting roles in *P. brassicae* response; however, this cannot be ruled out at this point and further research will be required to unravel this finding in full.

While many genes encoding TIR-NBS-LRR proteins, pathogenesis-related thaumatin superfamily proteins, and PR proteins have been found previously to be differentially expressed in different *Brassica* spp. with resistant or susceptible interactions **(Table 2**), and in seed-specific *AIL7* overexpressing siliques (Table 1; Singer et al., 2021), we did not observe similar, consistent changes in constitutive *AIL7* overexpression or T-DNA mutant roots in the current study (**Figure 7A**). Although this could be attributed to the different tissues assessed in each case, it is also possible that changes in the expression of these particular genes are not major factors contributing to the improved *P. brassicae* resistance observed in *AIL7* overexpression and mutant lines. Alternatively, phytohormones such as auxins, SA and JA have been shown previously to play a role in clubroot development (Jayasinghege et al., 2023), and AIL7, along with other family members, have been suggested to contribute to auxin biosynthesis and regulation (Krizek et al., 2020; Pinon et al., 2013). The seed-specific overexpression of *AIL7* was also previously found to lead to the altered expression of genes involved in auxin-, SA-and JA-related pathways (Singer et al., 2021), and as such, it is feasible that the role of *AIL7* in clubroot resistance could be linked to phytohormone biosynthesis and modification. In the current study, we did not observe any changes in auxin levels in either pathogen-inoculated or non-inoculated *AIL7* overexpression or mutant plants **(Figure S2)**, which suggests that *AIL7*-related alterations in clubroot tolerance in our transgenic lines were not auxin-dependent.

Conversely, in the absence of pathogen inoculation, we observed increased SA content and the upregulation of several major SA pathway genes (*ICS2*, *SID1, UGT74F2*) in both *AIL7* overexpression and knockout plants (**Figure 7B**). Since the accumulation of SA enables plants to initiate SAR and promotes broad-spectrum resistance, these results suggest that there may be some constitutive level of SAR present in *AIL7* overexpression and mutant lines, which is similar to what has been reported previously in *ail6ant* mutants (Krizek et al., 2016). In addition, the differential expression of *PR1* and *PR2* in the mutant lines (**Figure 7A**) further suggests activation of SAR, as PR protein production and accumulation are key features of SA-dependent SAR (Joshi et al., 2021). Furthermore, we observed a trend for reduced expression of *UGT76B1*, which coordinates the glucosylation of the group of defense activators (SA, isoleucic, N-hydroxy pipecolic acid) and leads to a SAR-like enhanced immune status when knocked out (Bauer et al., 2021), in *AIL7* overexpression and mutant lines compared to wild-type plants **(Figure 7B)**.

While higher endogenous SA accumulation or exogenous SA application is known to cause deleterious effects on plant growth (Dempsey & Klessig, 2017; Li et al., 2022; Pasternak et al., 2019), we did not notice any growth retardation in *AIL7* overexpression or mutant lines in the current study, and instead, improved growth of *AIL7* overexpression plants was observed **(Figures 2, 3 and 4)**. Since the detrimental impact of high levels of SA on plant growth is concentration-depend (Pasternak et al., 2019), it is possible that the levels of SA present in our transgenic lines were not elevated enough to induce such effects. On the other hand, increased transcript levels of *UGT74F2*, which is involved in SA glucosylation and controls the levels of unconjugated SA, were observed in *AIL7*-modulated lines in the absence of the pathogen, which may reduce the negative consequences of free SA (Dempsey & Klessig, 2017; Hou & Tsuda, 2022). Thus, our *AIL7* overexpression and mutant lines maintain a balance between defense response and growth, which is an important factor when developing resistant crop varieties.

While the maintenance of high SA levels in *P. brassicae* resistant/partially resistant cultivars has been reported in other studies (Jayasinghege et al., 2023; Lemarié et al., 2015; Prerostova et al., 2018), differences in JA levels among cultivars can be highly variable and may be affected by the host, pathotype, and disease development stage (Jayasinghege et al., 2023; Lemarié et al., 2015; Prerostova et al., 2018). While antagonistic behavior has typically been reported for SA and JA-based defense responses, as they are involved in biotrophic and necrotrophic interactions, respectively, synergistic interactions have also been observed in certain instances (Hou & Tsuda, 2022; Krizek et al., 2016; Ullah et al., 2022). Such a phenomenon would correspond with the increase in both SA and JA levels in AIL7-modulated lines in the absence of the pathogen in this study (**Figures 6A and 6B**). Intriguingly, a similar synergistic interaction between SA and JA was previously noted for *antail6* mutants, where greater SA and JA content in double mutants relative to a wild-type control was linked to enhanced resistance to *P. syringae* (Krizek et al., 2016). Alternatively, although there was a greater amount of JA in *AIL7* overexpression and mutant lines, as well as a corresponding upregulation of JA biosynthetic genes (*LOX* and *AOC*), it is possible that JA response could be suppressed by alterations in SA signaling, as described previously (Hou and Tsuda, 2022). In line with this, *JAZ1*, *8,* and *10* genes, which function as repressors of JA signaling (Liu & Timko, 2021), were upregulated in both *AIL7* overexpression and mutant lines following mock treatment. This type of interaction between phytohormones may enable plants to fine-tune their responses to various types of stress, thereby maintaining plant growth at optimal levels, even under limiting conditions (Beyer et al., 2021; Cui et al., 2018; Hou & Tsuda, 2022).

Interestingly, the levels of SA and JA did not show a similar trend in *P. brassicae*-inoculated *AIL7* overexpression and mutant lines compared to wild-type (**Figure 6B**), suggesting that there might be additional interactions, cross-talk, or regulatory mechanisms following infection. Indeed, while SA content was somewhat higher in *AIL7* overexpression lines compared to wild-type plants following inoculation, JA content was significantly enhanced in inoculated mutant lines. In the case of SA, a reduction in *UGT74F1*, *UGT74F2* and *UGT76B1* expression in *AIL7* overexpression lines post-inoculation (**Figure 7D**) might have contributed to increased levels of free SA content via the conversion of glycosylated SA to free SA **(Figure 6A)**. However, the downregulation of *SID2* in inoculated *AIL7* overexpression and mutant lines compared to wild-type, along with the reduced expression of *PR2* in overexpression lines, suggests that SA biosynthesis might be reduced in these lines compared to wild-type following inoculation. Since *AIL7*-modulated plants already have greater SA contents in the absence of the pathogen **(Figure 6A)**, further increases may be deleterious.

Increased JA levels and responses in the later stages of clubroot development have been reported previously in *P. brassicae-*resistant/partially resistant hosts (Jayasinghege et al., 2023; Lemarié et al., 2015). Interestingly, JA, ethylene, and brassinosteroid signaling have been found to be more important than SA signaling in terms of host response during the later stages of an incompatible interaction (Fu et al., 2019), and there is a tendency for SA to decline and JA to increase in the later stages of gall formation (Prerostova et al., 2018). Even though we did not observe significant changes in the expression of most JA biosynthetic genes in inoculated *AIL7*-modulated plants, the transcript levels of JA-responsive genes (i.e., *JAZ*) were significantly downregulated following inoculation in *ail7* mutant plants compared to wild-type (**Figure 7D**), which contrasted with uninoculated plants and suggests enhanced JA response in these lines. While *JAZ* expression in inoculated *AIL7* overexpressing lines demonstrated a similar trend the difference from wild-type plants was not significant (**Figure 7D**), which implies that the regulation of these phytohormonal pathways might be somewhat different between overexpressing and mutant lines under pathogen pressure. In line with this, we observed significantly enhanced JA content in *AIL7* mutant lines, whereas overexpressing plants displayed only a moderate, and insignificant, enhancement in JA content (**Figure 6B**). It is also important to note that *AIL7* overexpressing plants had significantly higher JA levels than wild-type without pathogen pressure, and further increases may have been deleterious as was suggested in the case of SA **(Figure 6A)**. Since we did not observe an increase in SA content in inoculated *AIL7*-modulated plants compared to wild-type, it is possible that upregulation in JA signaling could have contributed to a reduction in SA biosynthesis (Hou & Tsuda, 2022). A similar effect has been observed previously in *sid1* mutants, which are defective in SA accumulation and exhibit improved tolerance to *P. brassicae* (Lemarié et al., 2015).

Based on the accumulation of data from current and past studies, we have developed a tentative model to describe the role of this AP2/ERF family transcription factor in plant immunity against clubroot, one of the most severe soilborne disease of cruciferous plants (**Figure 9**). Our findings suggest that AIL7 plays a role by maintaining constitutive SAR even in the absence of *P. brassicae* infection. Such an effect involves both SA- and JA-related pathways, and this, rather than an accumulation of SA following *P. brassicae* challenge, primes the plants for improved resistance. This scenario is consistent with previous reports whereby constitutively elevated levels of SA were present in clubroot-resistant plants (Mencia et al., 2022; Prerostova et al., 2018). Moreover, we also observed a tendency towards a reduction in SA levels during disease development, as opposed to the maintenance of higher SA content throughout pathogenesis. This suggests that a fine-tuning of the balance between JA and SA is critical for determining precise defense mechanisms in response to *P. brassicae* inoculation in Arabidopsis. This involvement in SA and JA biosynthesis/ regulatory pathways could occur through direct interactions or the activation of key regulatory genes (Ruan et al., 2019). Such interactions might explain the synergistic accumulation of SA and JA that occurs in both inoculated and non-inoculated plants. Moreover, *AIL7* was shown previously to influence lipid biosynthesis and metabolism, impacting both storage and membrane lipid biosynthesis (Singer et al., 2021), and its close relative, *AIL6*, has also been demonstrated to be involved in storage and membrane lipid biosynthesis and metabolism (Kim et al., 2024; Liu et al., 2023; Zhang et al., 2016). This hints at the possibility that alterations in the expression of *AIL7* could feasibly lead to membrane lipid remodeling, which could trigger defense pathways such as the MAPK cascade, and phospholipid-based signaling cascade further activating phytohormone signaling (Mehta et al., 2021; Takasato et al., 2023). Further investigation into changes in membrane lipidome and targeted gene expression studies focusing on genes involved in lipid-mediated signaling cascades in plants with modified *AIL7* expression could further elucidate the relationships between clubroot resistance and AIL7 activity. In addition, interactions within this network are likely highly context-dependent, influenced by factors such as the type of stress, the stage of plant development, and environmental conditions. The further evaluation of these unanswered questions, along with an investigation of the specific gene targets of AIL7 and other proteins that could interact with AIL7 in multi-protein-complexes, will facilitate the elucidation of precise mechanisms behind AIL7-mediated clubroot resistance, and enable an assessment of the potential value of this gene as a target for developing clubroot-resistant *Brassica* crop cultivars downstream.

**Figure 9.**
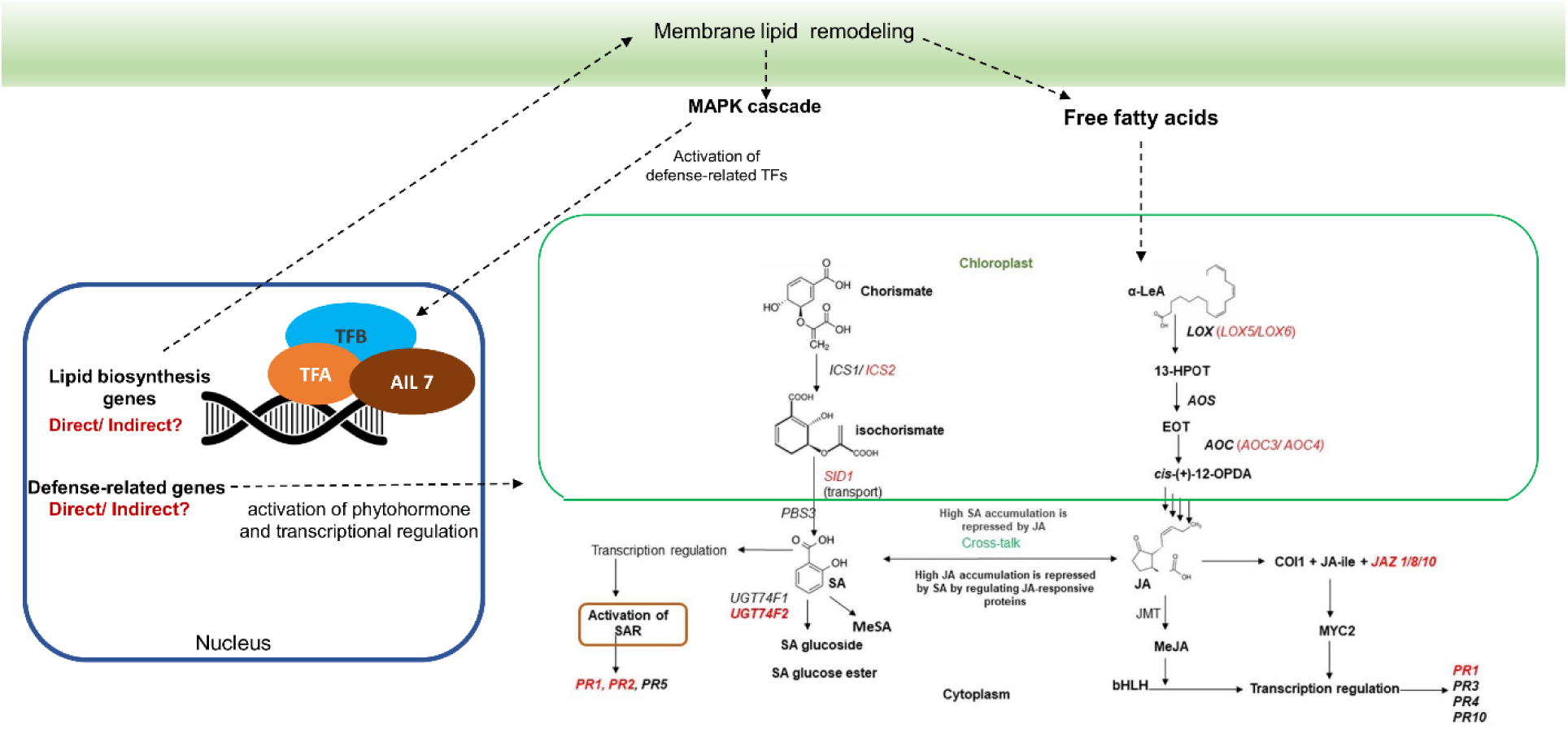
A tentative working model of the role of AIL7 in clubroot resistance. The possible direct and indirect effects of AIL7 on clubroot resistance are indicated. Red text indicates genes that are up/down-regulated in the current study. TF, transcription factor; TIR-NBS-LRR, Toll/interleukin-1 receptor-nucleotide binding site-leucine-rich repeat; MAPK, Mitogen-activated protein kinase; *ICS*, isochorismate synthase; *SID1*, salicylic acid induction deficient 1; *PR*, pathogenesis-related *UGT*, *UDP-DEPENDENT GLYCOSYLTRANSFERASE*; α-LeA, alpha-linolenic acid; *JAZ*, *JASMONATE-ZIM-DOMAIN*; *AOC*, *ALLENE OXIDE CYCLASE*; *LOX*s, *LIPOXYGENASE*; and *AOS, ALLENE OXIDE SYNTHASE*.

## Material and methods

### Plant material, constructs, and growth conditions

The *AIL7* (At5g65510) constitutive overexpression construct was generated by first amplifying the full-length 1497-nt *AIL7* coding region from cDNA derived from developing siliques of Arabidopsis using primers AtAIL7F1AgeI and AtAIL7R1BamHI (Table S1). These primers contain restriction sites to facilitate downstream cloning (Singer et al., 2021). The cloned fragment was inserted between the *Nicotiana tabacum* (tobacco) *tCUP3* constitutive promoter, and *Pisum sativum Ribulose-1,5-bisphosphate carboxylase* transcriptional terminator (rbcS-t) in a pGreen 0229 background (Hellens et al., 2000; Wickramarathna et al., 2015). Sequencing was carried out at every step of plasmid construction to ensure the correct identity of the resulting plasmid. The construct was then co-transformed into *Agrobacterium tumefaciens* GV3101 with the pSoup helper plasmid via electroporation (Hellens et al., 2000). *Agrobacterium*-mediated transformation of Col-0 Arabidopsis was conducted using the floral dip method (Clough & Bent, 1998).

T_1_ seeds were screened on selective medium containing phosphinothricin (25 mg/ L), and resistant seedlings were transferred to Sunshine LA4 potting mix (Sun Gro Horticulture, Vancouver, Canada) and grown to the next generation. Single-copy transgenic homozygous lines were identified through segregation analysis of T_2_ and T_3_ seeds as described previously (Jayawardhane et al., 2020; Singer et al., 2021). Arabidopsis *ail7* T-DNA insertion lines (SAIL 1167_C10, SAIL_267_G12) in the Col background were obtained from the Arabidopsis Biological Resource Center (http://arabidopsis.org), and homozygous lines for each mutant were identified using gene-specific primers and T-DNA border-specific primers as described previously (Krizek, 2009).

All *AIL7* overexpression (OE2, OE5, and OE12) and mutant lines (M1 and M2) selected for further assessment in this study, as well as the wild-type Col-0, were grown from seed in a growth chamber for 12 days at 22 °C/ 18 °C (day/night) with a photoperiod of 16 h day/8 h night and a light intensity of 250 μmolm^-2^s^-1^ prior to pathogen inoculation. Seeds were grown in 72 well (9*8) inserts filled with the previously mentioned potting mix. Two days prior to inoculation, the plants were moved to a greenhouse maintained at 24 °C/ 16 °C with a photoperiod of 16 h day/ 8 h night and a light intensity of 500 μmolm^-2^s^-1^. The plants were kept in the greenhouse until the completion of disease ratings and harvesting for other analyses.

### Assessment of plant and seed morphology of AIL7 overexpression and mutant lines

All morphological analyses of plants grown in potting mix were carried out using three independent overexpressing homozygous lines and two mutant lines (10-12 plants per construct), which were measured to estimate mean values for the following traits: seed size, total seed weight/plant (seed yield), seed weight/100 seeds, plant height, rosette diameter, and plant biomass. In each plant, rosette diameter and plant biomass were recorded 25 days after germination (before bolting) and plant height was measured 28 days after bolting. For seed size and seed weight analyses, plants were harvested at maturity and then cleaned using a sieve. Approximately 150-250 seeds from each line were photographed, and the images were used to quantify seed number using the particle counter function of QuPath v0.5.1 software. Seeds were weighed using an analytical balance (OHAUS Corporation, Florham Park, NJ, USA), and seed size was analyzed using ImageJ 1.53e software with the particle analysis function (Thresholding).

Root growth evaluation was carried out with minor adjustments based on previously established methods (Jayawardhane et al., 2020). In brief, surface-sterilized homozygous seeds were sown on half-strength MS medium in square plates (120 mm × 120 mm) without any antibiotics. These plates were cold stratified at 4 °C for 3 days, and were then oriented vertically inside a plant growth chamber with previously mentioned growth conditions. Measurements of root length were obtained 10 days post-germination, through the analysis of images using QuPath v0.5.1 software. This procedure was replicated three times, utilizing three separate seed batches in each case. To measure the length of hypocotyls in darkness, seeds underwent the same initial preparation and, following cold stratification, were briefly exposed to fluorescent white light (120 μM/m^2^/s) for 2 h to ensure synchronized germination. Following this treatment, the plates were covered in three layers of aluminum foil and positioned vertically in the growth chamber under the same conditions as described above, with the exception that light was excluded. Measurements of hypocotyl lengths were taken 10 days after the seeds had germinated.

### Pathogen material and inoculation

*P. brassicae* single-spore isolate 3H, classified based on the Canadian Clubroot Differential set (Strelkov et al., 2018), was used as the inoculum in this study due to the fact that it is widespread throughout canola-producing areas of Western Canada and exhibits high virulence against canola varieties lacking clubroot resistance (Hollman et al., 2021; Strelkov et al., 2018). Inoculum preparation and inoculation were carried out as described previously (Zhou et al., 2022). In brief, 5 g of frozen root galls, infected with pathotype 3H, were blended in 100 mL water and strained through eight layers of cheesecloth to remove plant tissue and other debris. The concentration of resting spores was determined using a hemocytometer and adjusted to a final concentration of 1.0 × 10^5^ resting spores per milliliter of water. Two-week old seedlings were inoculated according to a previously described method (Ludwig-Müller et al., 2017), whereby 2 mL of inoculum was applied to the potting mix adjacent to the base of each plant. Mock-inoculated control plants received an equivalent volume of water without resting spores following the same procedure. The experiments were also repeated with a higher concentration of inoculum (1× 10^6^ resting spores/mL water) to further assess the reaction of *AIL7* overexpression and T-DNA mutant lines.

### Disease rating

Clubroot severity was assessed 3 weeks following inoculation on a 0 – 3 scale based on Zhou et al. (2022), where: 0 = no galls, healthy root system; 1 = small galls, primarily on the lateral roots; 2 = small to medium galls on both the primary and lateral roots, moderate reduction in root system size; and 3 = large galls on the primary root, substantial reduction in root system size. The severity ratings were then used to calculate a DI for each replicate using a formula described previously (Strelkov et al., 2006): DI (%) = [(n_1_ × 1 + n_2_ × 2 + n_3_ × 3)/(N × 3)] × 100, where n_1_, n_2_ and n_3_ are the number of plants in each severity class and N is the total number of plants tested. The experiment was replicated four times with 24 plants per Arabidopsis line (OE2, OE5, OE12, M1, M2, and wild-type) in each replicate (144 plants per replicate), in a completely randomized design.

### Hormone Profiling

SA, JA, and auxin levels were measured as described previously (Müller et al., 2002), with minor modifications. Four biological replicates were used in the case of mutants and overexpression lines, while three biological replicates were included for the wild-type control lines. Each replicate consisted of 15-20 plants and a similar number of replicates and plants were inoculated with water as the mock test. Root samples were harvested 14 days post-inoculation, washed thoroughly with distilled water, blot-dried, flash-frozen in liquid nitrogen, and stored at - 80℃.

The measurement of phytohormones was conducted using LC-MRM/MS by a service provider (University of Victoria - Genome BC Proteomics Centre, Victoria BC, Canada). In brief, each sample was pulverized to a fine paste at 30 Hz for 3 min in a 5 mL tissue homogenization tube on a MM 400 mixer mill with the aid of two metal balls. A 50 mg aliquot of each sample was weighed into a 2 mL tissue homogenization tube, and 1 mL of 80% aqueous methanol was added to each tube. The samples were homogenized for metabolite extraction at 30 Hz for 3 min, followed by ultra-sonication in an ice-water bath for 10 min, and subsequent centrifugal clarification at 21,000 g at 10 °C for 10 min. A 100 µL aliquot of the clear supernatant of each sample was mixed with an equal volume of an internal standard containing indole 3-acetic acid- d2 and salicylic acid-d4 (0.1 µM of each). Ten serially diluted calibration solutions containing standards of the measured compounds at a concentration range of 0.0001 to 20 µM were also prepared in 80% aqueous methanol. One hundred µL of each calibration solution was mixed with an equal volume of the same internal standard solution, and 10 μL aliquots of all resulting solutions were injected onto a C18 column (2.1*150 mm, 1.8 µm) to run LC-MRM/MS with (-) ion detection on an Agilent 1290 UHPLC system coupled to an Agilent 6495C triple-quadrupole mass spectrometer. The mobile phase consisted of 2 mM ammonium acetate solution (A) and acetonitrile (B) for binary solvent gradient elution under an optimized condition. Linear regression curves of the individual compounds were generated with the data acquired from the calibration solutions. The concentration of each compound detected in the samples was calculated with internal standard calibration by interpolating the calibration curves with the data acquired from the sample solutions, with an appropriate concentration range for each compound. For the compounds without isotope-labeled internal standards, salicylic acid-d4 was used as a common internal standard.

### Total RNA extraction, cDNA synthesis and quantitative PCR

Total RNA was extracted using the Spectrum™ Plant Total RNA Kit with the associated On-Column DNase I Digest Set (Sigma-Aldrich) to remove trace DNA. cDNA synthesis was carried out using the Superscript IV First-Strand cDNA Synthesis Kit according to the manufacturer’s instructions (Invitrogen, Life Technologies Inc.), with 350 ng total RNA as template and an oligo dT primer. All qPCR assays were conducted using a Step OnePlus Real-Time PCR System (Life Technologies) in a total volume of 10 µl with SYBR Green PCR Master Mix (Molecular Biology Service Unit, University of Alberta, Edmonton, AB) as described in a previous study (Neil et al., 2024). The following thermal parameters were used for amplification: 95 °C for 2 min, followed by 40 cycles of 95 °C for 15 s and 60 ^0^C for 1 min. Dissociation curves were generated for each reaction to confirm amplification specificity. Levels of gene expression relative to the internal control were obtained using the comparative Ct method (Schmittgen & Livak, 2008).

To confirm *AIL7* overexpression, total RNA was extracted from the roots of 28-day old homozygous *AIL7* overexpression lines and control wild-type Col-0 lines. *AIL7* gene-specific primers (AIL7qPCRF and AIL7qPCRR) were utilized to amplify a region of *AIL7* while primers PP2AA3F and PP2AA3R were used to amplify a region of the internal control gene *PROTEIN PHOSPHATASE 2A SUBUNIT 3* (*PP2AA3*) (Table S1; Singer et al., 2016, 2021). In addition, the expression levels of *AIL3*, *AIL4*, *AIL5*, and *AIL6*, which are other family members which may have partial redundancy with *AIL7*, were also analyzed in root and leaf tissues. qPCR assays were performed using three biological replicates and three technical replicates, with 1 µl of 1/10 diluted cDNA as a template.

To analyze the expression levels of defense- and hormone-related genes, total RNA was extracted from the roots of both *P. brassicae*-inoculated and mock-inoculated plants (14 days post-inoculation). qPCR assays were performed using 3-4 biological replicates, which represented approximately 20 individual plants per sample (OE, mutants or WT), and three technical replicates, with 1 µl of 1/5 diluted cDNA as a template and gene-specific primers listed in Table S1.

### Statistical analyses

For all analyses, two-tailed student’s t-tests were conducted and the data means represent the average of biological replicates. Differences were considered significant at *p* ≤ 0.05.

## Supplementary files

Figure S1. Rating scale and the evaluation of *AIL7* overexpression, mutant, and wild-type (Col-0) Arabidopsis lines following inoculation with *Plasmodiophora brassicae* pathotype 3H (1*10^6^ spores/ mL).

Figure S2. Auxin (IAA, IBA and IPA) measurements in *P. brassicae* inoculated and non-inoculated Arabidopsis roots.

Table S1. Primers used in this study.

Table S2. List of DEGs with putative importance in disease resistance, identified by mining previous comparative transcriptomic data between seed-specific *AIL7* Arabidopsis overexpression and wild-type siliques (Singer et al. 2021). Green indicates the upregulation of genes compared to wild-type while red indicates downregulation

## Abbreviations used

TF: transcription factor
AP2/ERF: Apetala2/Ethylene Responsive Factor
AIL: *AINTEGUMENTA LIKE*
JA: jasmonic acid
SA: Salicylic acid
PR: *PATHOGENESIS-RELATED*
*SID*: *SALICYLIC ACID INDUCTION DEFICIENT*
NBS-LRR: nucleotides binding-site-leucine-rich repeat
ETI: effector-triggered immune
*UGT*: *UDP-DEPENDENT GLYCOSYLTRANSFERASE*
DAI: Days after inoculation
SAR: systemic acquired resistance
ROS: reactive oxygen species
MAPK: mitogen-activated protein kinase
DI: Disease index

## Acknowledgment

We thank Drs. Jun Han and David Schibli in Uvic-Genome BC Proteomics Centre, Victoria, for their technical support in the hormone analysis. Furthermore, we gratefully acknowledge the technical support of Dr. Bin Shan, Siyu Wang, Brock Mason, and Solomiya Kucharyshyn in plant maintenance and sample collection, and Dr. Charitha Jayasinghege and Qinqin Zhou for their valuable discussion. The research was supported by Alberta Canola Producers Commission (GC, SDS, SES and SFH), Results Driven Agriculture Research (GC, SDS, SES and SFH), Western Grains Research Foundation (GC, SDS, SES and SFH), the University of Alberta Start-up Research Grant (GC), the Natural Science and Engineering Research Council of Canada (NSERC) Discovery Grants (GC and SES), the Canada Research Chairs Program (GC), Canada Foundation for Innovation-John R. Evans Leaders Fund (Project number 41867) and Research Capacity Program of Alberta (RCP-22-023-SEG).

## Author contribution statement

KNJ, GC and SDS, conceived and designed the experiments. KNJ, TS and VPM carried out all phenotypic and molecular experiments. SES provided all the pathogen material and other resources relevant to clubroot analysis. KNJ carried out all the data analysis and wrote the first draft. GC, SDS, SES and SFH supervised the research and acquired funding. All authors revised the manuscript.

## References

Adhikary, D., Kisiala, A., Sarkar, A., Basu, U., Rahman, H., Emery, N., & Kav, N. N. V. (2022). Early-stage responses to *Plasmodiophora brassicae* at the transcriptome and metabolome levels in clubroot resistant and susceptible oilseed *Brassica napus*. Molecular Omics, 18(10), 991– 1014. 10.1039/d2mo00251e

Aida, M., Beis, D., Heidstra, R., Willemsen, V., Blilou, I., Galinha, C., Nussaume, L., Noh, Y. S., Amasino, R., & Scheres, B. (2004). The *PLETHORA* genes mediate patterning of the Arabidopsis root stem cell niche. Cell, 119(1), 109–120. 10.1016/j.cell.2004.09.018

Bao, H., Ding, Y., Yang, F., Zhang, J., Xie, J., Zhao, C., Du, K., Zeng, Y., Zhao, K., Li, Z., & Yang, Z. (2022). Gene silencing, knockout and over-expression of a transcription factor *ABORTED MICROSPORES* (*SlAMS*) strongly affects pollen viability in tomato (*Solanum lycopersicum*). BMC Genomics, 23, 346. 10.1186/s12864-022-08549-x

Bauer, S., Mekonnen, D. W., Hartmann, M., Yildiz, I., Janowski, R., Lange, B., Geist, B., Zeier, J., & Schäffner, A. R. (2021). UGT76B1, a promiscuous hub of small molecule-based immune signaling, glucosylates N-hydroxypipecolic acid, and balances plant immunity. Plant Cell, 33(3), 714–734. 10.1093/plcell/koaa044

Beyer, S. F., Bel, P. S., Flors, V., Schultheiss, H., Conrath, U., & Langenbach, C. J. G. (2021). Disclosure of salicylic acid and jasmonic acid-responsive genes provides a molecular tool for deciphering stress responses in soybean. Scientific Reports, 11(1): 20600. 10.1038/s41598-021-00209-6

Blümke, A., Somerville, S. C., & Voigt, C. A. (2013). Transient expression of the *Arabidopsis thaliana* callose synthase *PMR4* increases penetration resistance to powdery mildew in barley. Advances in Bioscience and Biotechnology, 04(08), 810–813. 10.4236/abb.2013.48106

Boutilier, K., Offringa, R., Sharma, V. K., Kieft, H., Ouellet, T., Zhang, L., Hattori, J., Liu, C. M., Van Lammeren, A. A. M., Miki, B. L. A., Custers, J. B. M., & Van Lookeren Campagne, M. M. (2002). Ectopic expression of *BABY BOOM* triggers a conversion from vegetative to embryonic growth. Plant Cell, 14(8), 1737–1749. 10.1105/tpc.001941

Chen, J., Pang, W., Chen, B., Zhang, C., & Piao, Z. (2016). Transcriptome analysis of *Brassica rapa* near-isogenic lines carrying clubroot-resistant and -susceptible alleles in response to *Plasmodiophora brassicae* during early infection. Frontiers in Plant Science, 6, 1183. 10.3389/fpls.2015.01183

Clough, S. J., & Bent, A. F. (1998). Floral dip: A simplified method for Agrobacterium-mediated transformation of *Arabidopsis thaliana*. Plant Journal, 16(6), 735–743. 10.1046/j.1365-313X.1998.00343.x

Cui, H., Qiu, J., Zhou, Y., Bhandari, D. D., Zhao, C., Bautor, J., Parker., J.E. (2018). Antagonism of transcription factor MYC2 by EDS1/PAD4 complexes bolsters salicylic acid defense in Arabidopsis effector-triggered immunity. Molecular Plant 11, 1053–1066. 10.1016/j.molp.2018.05.007

Cusack, S. A., Wang, P., Lotreck, S. G., Moore, B. M., Meng, F., Conner, J. K., Krysan, P. J., Lehti-Shiu, M. D., & Shiu, S. H. (2021). Predictive models of genetic redundancy in *Arabidopsis thaliana*. Molecular Biology and Evolution, 38(8), 3397–3414. 10.1093/molbev/msab111

Delgado-Baquerizo, M., Guerra, C. A., Cano-Díaz, C., Egidi, E., Wang, J. T., Eisenhauer, N., Singh, B. K., & Maestre, F. T. (2020). The proportion of soil-borne pathogens increases with warming at the global scale. Nature Climate Change, 10(6), 550–554. 10.1038/s41558-020-0759-3

Dempsey, D. A., & Klessig, D. F. (2017). How does the multifaceted plant hormone salicylic acid combat disease in plants and are similar mechanisms utilized in humans? BMC Biology, 15, 23. 10.1186/s12915-017-0364-8

Ding, J., Ruan, C., Guan, Y., & Krishna, P. (2018). Identification of microRNAs involved in lipid biosynthesis and seed size in developing sea buckthorn seeds using high-throughput sequencing. Scientific Reports, 8(1), 1–15. 10.1038/s41598-018-22464-w

Ding, Q., Cui, B., Li, J., Li, H., Zhang, Y., Lv, X., Qiu, N., Liu, L., Wang, F., & Gao, J. (2018). Ectopic expression of a *Brassica rapa AINTEGUMENTA* gene (*BrANT-1*) increases organ size and stomatal density in Arabidopsis. Scientific Reports, 8(1), 1–13. 10.1038/s41598-018-28606-4

Dixon G.R. (2009). *Plasmodiophora brassicae* in its environment. Journal of Plant Growth Regulation, 28, 212–228. 10.1007/s00344-009-9098-3

Dodds, P. N., Chen, J., & Outram, M. A. (2024). Pathogen perception and signaling in plant immunity. The Plant Cell, 36(5), 1465–1481. 10.1093/plcell/koae020

Dong, L., Cheng, Y., Wu, J., Cheng, Q., Li, W., Fan, S., Jiang, L., Xu, Z., Kong, F., Zhang, D., Xu, P., & Zhang, S. (2015). Overexpression of *GmERF5*, a new member of the soybean EAR motif-containing ERF transcription factor, enhances resistance to *Phytophthora sojae* in soybean. Journal of Experimental Botany, 66(9), 2635–2647. 10.1093/jxb/erv078

Dos Santos, C., & Franco, O. L. (2023). Pathogenesis-related proteins (PRs) with enzyme activity activating plant defense responses. Plants, 12(11), 2226. 10.3390/plants12112226

Feng, K., Hou, X. L., Xing, G. M., Liu, J. X., Duan, A. Q., Xu, Z. S., Li, M. Y., Zhuang, J., & Xiong, A. S. (2020). Advances in AP2/ERF super-family transcription factors in plant. Critical Reviews in Biotechnology, 40(6), 750–776. 10.1080/07388551.2020.1768509

Fu, P., Piao, Y., Zhan, Z., Zhao, Y., Pang, W., Li, X., & Piao, Z. (2019). Transcriptome profile of *Brassica rapa* L. reveals the involvement of jasmonic acid, ethylene, and brassinosteroid signaling pathways in clubroot resistance. Agronomy, 9(10), 589. 10.3390/agronomy9100589

Gan, C., Yan, C., Pang, W., Cui, L., Fu, P., Yu, X., Qiu, Z., Zhu, M., Piao, Z., & Deng, X. (2022). Identification of novel locus RsCr6 related to clubroot resistance in radish (*Raphanus sativus* L.). Frontiers in Plant Science, 13, 866211. 10.3389/fpls.2022.866211

Han, H., & Krizek, B. A. (2016). AINTEGUMENTA-LIKE6 can functionally replace AINTEGUMENTA but alters Arabidopsis flower development when misexpressed at high levels. Plant Molecular Biology, 92(4–5), 597–612. 10.1007/s11103-016-0535

Hasan, J., Megha, S., & Rahman, H. (2021). Clubroot in brassica: Recent advances in genomics, breeding, and disease management. Genome, 64(8), 735–760. 10.1139/gen-2020-0089

He, P., Warren, R. F., Zhao, T., Shan, L., Zhu, L., Tang, X., & Zhou, J.-M. (2001). Overexpression of *Pti5* in tomato potentiates pathogen-induced defense gene expression and enhances disease resistance to *Pseudomonas syringae pv*. Tomato. The American Phytopathological Society, 14(12), 1453–1457. 10.1094/MPMI.2001.14.12.1453

Hellens, R., Edwards, E., Leyland, N., Bean, S., & Mullineaux, P. (2000). pGreen: a versatile and flexible binary Ti vector for Agrobacterium-mediated plant transformation. Plant Molecular Biology, 42(6), 819–832. 10.1023/A:1006496308160

Hofhuis, H., Laskowski, M., Du, Y., Prasad, K., Grigg, S., Pinon, V., & Scheres, B. (2013). Phyllotaxis and rhizotaxis in Arabidopsis are modified by three plethora transcription factors. Current Biology, 23(11), 956–962. 10.1016/j.cub.2013.04.048

Hollman, K. B., Hwang, S. F., Manolii, V. P., & Strelkov, S. E. (2021). Pathotypes of *Plasmodiophora brassicae* collected from clubroot resistant canola (*Brassica napus* L.) cultivars in western Canada in 2017-2018. Canadian Journal of Plant Pathology, 43(4), 622–630. 10.1080/07060661.2020.1851893

Horstman, A., Willemsen, V., Boutilier, K., & Heidstra, R. (2016). AINTEGUMENTA-LIKE proteins : hubs in a plethora of networks. Trends in Plant Science, 19(3), 146–157. 10.1016/j.tplants.2013.10.010

Hou, S., & Tsuda, K. (2022). Salicylic acid and jasmonic acid crosstalk in plant immunity. Essays in biochemistry, 66(5), 647–656. 10.1042/EBC20210090

Irani, S., Trost, B., Waldner, M., Nayidu, N., Tu, J., Kusalik, A. J., Todd, C. D., Wei, Y., & Bonham-Smith, P. C. (2018). Transcriptome analysis of response to *Plasmodiophora brassicae* infection in the Arabidopsis shoot and root. BMC Genomics, 19(1), 23. 10.1186/s12864-017-4426-7

Jayasinghege, C. P. A., Ozga, J. A., Manolii, V. P., Hwang, S. F., & Strelkov, S. E. (2023). Impact of susceptibility on plant hormonal composition during clubroot disease development in canola (*Brassica napus*). Plants, 12(16), 2899. 10.3390/plants12162899

Jayawardhane, K. N., Singer, S. D., Ozga, J. A., Rizvi, S. M., Weselake, R. J., & Chen, G. (2020). Seed-specific down-regulation of Arabidopsis *CELLULOSE SYNTHASE 1* or *9* reduces seed cellulose content and differentially affects carbon partitioning. Plant Cell Reports, 39, 953– 969. doi: 10.1007/s00299-020-02541-z

Jia, H., Wei, X., Yang, Y., Yuan, Y., Wei, F., Zhao, Y., Yang, S., Yao, Q., Wang, Z., Tian, B., & Zhang, X. (2017). Root RNA-seq analysis reveals a distinct transcriptome landscape between clubroot-susceptible and clubroot-resistant Chinese cabbage lines after *Plasmodiophora brassicae* infection. Plant and Soil, 421(1–2), 93–105. 10.1007/s11104-017-3432-5

Joshi, V., Joshib, N., Vyasc, A., Jadhav, S.K. (2021) Pathogenesis-related proteins: Role in plant defense. In Jogaiah, S. Ed., Biocontrol Agents and Secondary Metabolites, Woodhead Publishing, 573–590. 10.1016/B978-0-12-822919-4.00025-9

Khalid, M., Rahman, S.U., Kayani, S., Khan, A.A., Gul, H., & Hui, N. (2022). *Plasmodiophora brassicae*–The causal agent of clubroot and its biological control/suppression with fungi. South African Journal of Botany.147,325–331. 10.1016/j.sajb.2022.01.032

Kim, I., Do, H., Park, M. E., & Kim, H. U. (2024). Multiple transcription factors of *Arabidopsis thaliana* that are activated by LEAFY COTYLEDON 2 regulate triacylglycerol biosynthesis. Plant Journal. 10.1111/tpj.16762

Kopec, P. M., Mikolajczyk, K., Jajor, E., Perek, A., Nowakowska, J., Obermeier, C., Chawla, H. S., Korbas, M., Bartkowiak-Broda, I., & Karlowski, W. M. (2021). Local duplication of *TIR-NBS-LRR* gene marks clubroot resistance in *Brassica napus cv. Tosca*. Frontiers in Plant Science, 12, 639631. 10.3389/fpls.2021.639631

Krizek, B. A. (2009). *AINTEGUMENTA* and *AINTEGUMENTA-LIKE6* act redundantly to regulate Arabidopsis floral growth. Plant Physiology, 150, 1916–1929. 10.1104/pp.109.141119

Krizek, B. A. (2015). AINTEGUMENTA-LIKE genes have partly overlapping functions with AINTEGUMENTA but make distinct contributions to Arabidopsis thaliana flower development. Journal of Experimental Botany, 66(15), 4537–4549. 10.1093/jxb/erv224

Krizek, B. A., Bequette, C. J., Xu, K., Blakley, I. C., Fu, Z. Q., Stratmann, J. W., & Loraine, A. E. (2016). RNA-Seq links the transcription factors AINTEGUMENTA and AINTEGUMENTA-LIKE6 to cell wall remodeling and plant defense pathways. Plant Physiology, 171(3), 2069– 2084. 10.1104/pp.15.01625

Krizek, B. A., Blakley, I. C., Ho, Y. Y., Freese, N., & Loraine, A. E. (2020). The Arabidopsis transcription factor AINTEGUMENTA orchestrates patterning genes and auxin signaling in the establishment of floral growth and form. Plant Journal, 103(2),752–768. 10.1111/tpj.14769

Krizek, B. A., Iorio, C. B., Higgins, K., & Han, H. (2023). Differences in both expression and protein activity contribute to the distinct functions of AINTEGUMENTA compared with AINTEGUMENTA-LIKE 5 and AINTEGUMENTA-LIKE 7. Plant Molecular Biology, 113(1–3), 75–88. 10.1007/s11103-023-01374-0

Lemarié, S., Robert-Seilaniantz, A., Lariagon, C., Lemoine, J., Marnet, N., Jubault, M., Manzanares-Dauleux, M. J., & Gravot, A. (2015). Both the jasmonic acid and the salicylic acid pathways contribute to resistance to the biotrophic clubroot agent *Plasmodiophora brassicae* in Arabidopsis. Plant and Cell Physiology, 56(11), 2158–2168. 10.1093/pcp/pcv127

Li, A., Sun, X., & Liu, L. (2022). Action of salicylic acid on plant growth. Frontiers in Plant Science, 13, 878076. 10.3389/fpls.2022.878076

Licausi, F., Ohme-Takagi, M., & Perata, P. (2013). APETALA2/Ethylene responsive factor (AP2/ERF) transcription factors: mediators of stress responses and developmental programs. The New phytologist, 199(3), 639–649. 10.1111/nph.12291

Liu, H., & Timko, M. P. (2021). Jasmonic acid signaling and molecular crosstalk with other phytohormones. International Journal of Molecular Sciences, 22(6), 1–24. 10.3390/ijms22062914

Liu, X., Li, N., Chen, A., Saleem, N., Jia, Q., Zhao, C., Li, W., & Zhang, M. (2023). FUSCA3-induced AINTEGUMENTA-like 6 manages seed dormancy and lipid metabolism. Plant Physiology, 193(2), 1091–1108. 10.1093/plphys/kiad397

Ludwig-Müller, J., Auer, S., Jülke, S., & Marschollek, S. (2017). Manipulation of auxin and cytokinin balance during the *Plasmodiophora brassicae*–*Arabidopsis thaliana* interaction. Methods molecular Biology, 1569:41–60. 10.1007/978-1-4939-6831-2_3

Ludwig-Müller, J., Epstein, E., & Hilgenberg, W. (1996). Auxin-conjugate hydrolysis in Chinese cabbage: Characterization of an amidohydrolase and its role during infection with clubroot disease. Physiologia Plantarum, 97(4), 627–634. 10.1111/j.1399-3054.1996.tb00525.x

Ma, N., Sun, P., Li, Z. Y., Zhang, F. J., Wang, X. F., You, C. X., Zhang, C. L., & Zhang, Z. (2024). Plant disease resistance outputs regulated by AP2/ERF transcription factor family. Stress Biology, 4(1), 2. 10.1007/s44154-023-00140-y

Malley, R. C. O., Carol, S., Song, L., Galli, M., Ecker, J. R., Malley, R. C. O., Huang, S. C., Song, L., Lewsey, M. G., Bartlett, A., & Nery, J. R. (2016). Cistrome and epicistrome features shape the regulatory DNA landscape resource cistrome and epicistrome features shape the regulatory DNA landscape. Cell, 165(5), 1280–1292. 10.1016/j.cell.2016.04.038

Mehta, S., Chakraborty, A., Roy, A., Singh, I. K., & Singh, A. (2021). Fight hard or die trying: Current status of lipid signaling during plant–pathogen interaction. Plants, 10(6), 1098. 10.3390/plants10061098

Mencia, R., Welchen, E., Auer, S., & Ludwig-Müller, J. (2022). A novel target (Oxidation Resistant 2) in *Arabidopsis thaliana* to reduce clubroot disease symptoms via the salicylic acid pathway without growth penalties. Horticulturae, 8(1), 9. 10.3390/horticulturae8010009

Mudunkothge, J. S., & Krizek, B. A. (2012). Three Arabidopsis *AIL / PLT* genes act in combination to regulate shoot apical meristem function. The Plant Journal, 71, 108–121. 10.1111/j.1365-313X.2012.04975.x

Müller, A., Düchting, P., & Weiler, E. W. (2002). A multiplex GC-MS/MS technique for the sensitive and quantitative single-run analysis of acidic phytohormones and related compounds, and its application to *Arabidopsis thaliana*. Planta, 216(1), 44–56. 10.1007/s00425-002-0866-6

Neil, G. J., Kluttig, K. H., & Allison, W. T. (2024). Determining photoreceptor cell identity: Rod versus cone fate governed by tbx2b opposing nrl. Investigative Ophthalmology and Visual Science, 65(1), 39. 10.1167/iovs.65.1.39

Ng, D. W. K., Abeysinghe, J. K., & Kamali, M. (2018). Regulating the regulators: The control of transcription factors in plant defense signaling. International Journal of Molecular Sciences. 19(12), 3737. 10.3390/ijms19123737

Nie, S., & Wang, D. (2023). AP2/ERF transcription factors for tolerance to both biotic and abiotic stress factors in plants. Tropical Plant Biology. 16(3), 105–112. 10.1007/s12042-023-09339-9

Nishimura Marc T., Stein Monica, Hou Bi-Huei, Vogel John P., Edwards, H., & Somerville Shauna C. (2003). Loss of a callose synthase results in salicylic acid–dependent disease resistance. Science, 301(5635), 969–72. 10.1126/science.1086716

Nole-Wilson, S., Tranby, T. L., & Krizek, B. A. (2005). *AINTEGUMENTA-like* (*AIL*) genes are expressed in young tissues and may specify meristematic or division-competent states. Plant Molecular Biology, 57(5), 613–628. 10.1007/s11103-005-0955-6

Pasternak, T., Groot, E. P., Kazantsev, F. V., Teale, W., Omelyanchuk, N., Kovrizhnykh, V., Palme, K., & Mironova, V. V. (2019). Salicylic acid affects root meristem patterning via auxin distribution in a concentration-dependent manner. Plant Physiology, 180(3), 1725–1739. 10.1104/pp.19.00130

Peng, J. (2019). Gene redundancy and gene compensation: An updated view. Journal of Genetics and Genomics, 46(7), 329–333. 10.1016/j.jgg.2019.07.001

Peng, Y., Yang, J., Li, X., & Zhang, Y. (2021). Salicylic acid: Biosynthesis and signaling. Annual Review of Plant Biology, 17(72), 761–791. 10.1146/annurev-arplant-081320

Pinon, V., Prasad, K., Grigg, S. P., Sanchez-Perez, G. F., & Scheres, B. (2013). Local auxin biosynthesis regulation by PLETHORA transcription factors controls phyllotaxis in Arabidopsis. Proceedings of the National Academy of Sciences, 110(3), 1107–1112. 10.1073/pnas.1213497110

Prelich, G. (2012). Gene overexpression: Uses, mechanisms, and interpretation. Genetics, 190(3), 841–854. 10.1534/GENETICS.111.136911

Prerostova, S., Dobrev, P. I., Konradyova, V., Knirsch, V., Gaudinova, A., Kramna, B., Kazda, J., Ludwig-Müller, J., & Vankova, R. (2018). Hormonal responses to *plasmodiophora brassicae* infection in *brassica napus* cultivars differing in their pathogen resistance. International Journal of Molecular Sciences, 19(12), 4024. 10.3390/ijms19124024

Ruan, J., Zhou, Y., Zhou, M., Yan, J., Khurshid, M., Weng, W., Cheng, J., & Zhang, K. (2019). Jasmonic acid signaling pathway in plants. International Journal of Molecular Sciences, 20(10), 2479. 10.3390/ijms20102479

Santillán Martínez, M. I., Bracuto, V., Koseoglou, E., Appiano, M., Jacobsen, E., Visser, R. G. F., Wolters, A. M. A., & Bai, Y. (2020). CRISPR/Cas9-targeted mutagenesis of the tomato susceptibility gene *PMR4* for resistance against powdery mildew. BMC Plant Biology, 20(1), 284. 10.1186/s12870-020-02497-y

Santuari, L., Sanchez-Perez, G. F., Luijten, M., Rutjens, B., Terpstra, I., Berke, L., Gorte, M., Prasad, K., Bao, D., Timmermans-Hereijgers, J. L. P. M., Maeo, K., Nakamura, K., Shimotohno, A., Pencik, A., Novak, O., Ljung, K., van Heesch, S., de Bruijn, E., Cuppen, E., Willemsen V, Mähönen A. P., Lukowitz W., Snel B., de Ridder D., Scheres B., Heidstra R. (2016). The *PLETHORA* gene regulatory network guides growth and cell differentiation in Arabidopsis roots. Plant Cell, 28(12), 2937–2951. 10.1105/tpc.16.00656

Schmittgen, T. D., & Livak, K. J. (2008). Analyzing real-time PCR data by the comparative CT method. Nature Protocols, 3(6), 1101–1108. 10.1038/nprot.2008.73

Singer, S. D., Chen, G., Mietkiewska, E., Tomasi, P., Jayawardhane, K., Dyer, J. M., & Weselake, R. J. (2016). Arabidopsis GPAT9 contributes to synthesis of intracellular glycerolipids but not surface lipids. Journal of Experimental Botany, 67(15), 4627–4638. 10.1093/jxb/erw242

Singer, S. D., Jayawardhane, K. N., Jiao, C., Weselake, R. J., & Chen, G. (2021). The effect of *AINTEGUMENTA-LIKE 7* over-expression on seed fatty acid biosynthesis, storage oil accumulation and the transcriptome in Arabidopsis thaliana. Plant Cell Reports, 40(9), 1647– 1663. 10.1007/s00299-021-02715-3

Strelkov, S. E., Hwang, S. F., Manolii, V. P., Cao, T., Fredua-Agyeman, R., Harding, M. W., Peng, G., Gossen, B. D., Mcdonald, M. R., & Feindel, D. (2018). Virulence and pathotype classification of *Plasmodiophora brassicae* populations collected from clubroot resistant canola (*Brassica napus*) in Canada. Canadian Journal of Plant Pathology, 40(2), 284–298. 10.1080/07060661.2018.1459851

Strelkov, S. E., Tewari, J. P., & Smith-Degenhardt, E. (2006). Characterization of *Plasmodiophora brassicae* populations from Alberta, Canada. Canadian Journal of Plant Pathology, 28(3), 467–474. 10.1080/07060660609507321

Takasato, S., Bando, T., Ohnishi, K., Tsuzuki, M., Hikichi, Y., & Kiba, A. (2023). Phosphatidylinositol-phospholipase C3 negatively regulates the hypersensitive response via complex signaling with MAP kinase, phytohormones, and reactive oxygen species in *Nicotiana benthamiana*. Journal of Experimental Botany, 74(15), 4721–4735. 10.1093/jxb/erad184

Tayengwa, R., Sharma Koirala, P., Pierce, C. F., Werner, B. E., & Neff, M. M. (2020). Overexpression of *AtAHL20* causes delayed flowering in Arabidopsis via repression of *FT* expression. BMC Plant Biology, 20(1). 559. 10.1186/s12870-020-02733-5

Ullah, C., Schmidt, A., Reichelt, M., Tsai, C. J., & Gershenzon, J. (2022). Lack of antagonism between salicylic acid and jasmonate signalling pathways in poplar. New Phytologist, 235(2), 701–717. 10.1111/nph.18148

Vogel, J., & Somerville, S. (1999). Isolation and characterization of powdery mildew-resistant Arabidopsis mutants. Proceedings of the National Academy of Sciences, 97(4), 1897–1902. 10.1073/pnas.03053199

Wang, X., Zhang, J., Zhang, J., Zhou, C., & Han, L. (2022). Genome-wide characterization of AINTEGUMENTA-LIKE family in *Medicago truncatula* reveals the significant roles of AINTEGUMENTAs in leaf growth. Frontiers in Plant Science, 13, 1050462. 10.3389/fpls.2022.1050462

Wang, Y., Pruitt, R. N., Nürnberger, T., & Wang, Y. (2022). Evasion of plant immunity by microbial pathogens. Nature Reviews Microbiology, 20(8), 449–464. 10.1038/s41579-022-00710-3

Wickramarathna, A. D., Siloto, R. M. P., Mietkiewska, E., Singer, S. D., Pan, X., & Weselake, R. J. (2015). Heterologous expression of flax *PHOSPHOLIPID: DIACYLGLYCEROL CHOLINEPHOSPHOTRANSFERASE* (*PDCT*) increases polyunsaturated fatty acid content in yeast and Arabidopsis seeds. BMC Biotechnology, 15(1), 1–15. 10.1186/s12896-015-0156-6

Yu, F., Zhang, X., Peng, G., Falk, K. C., Strelkov, S. E., & Gossen, B. D. (2017). Genotyping-by-sequencing reveals three QTL for clubroot resistance to six pathotypes of *Plasmodiophora brassicae* in *Brassica rapa*. Scientific Reports, 7(1), 4516. 10.1038/s41598-017-04903-2

Zahr, K., Yang, Y., Sarkes, A., Dijanovic, S., Fu, H., Harding, M. W., Feindel, D., & Feng, J. (2022). *Plasmodiophora brassicae* infection threshold—how many resting spores are required for generating clubroot galls on canola (*Brassica napus*). Journal of Plant Diseases and Protection, 129(2), 387–394. 10.1007/s41348-022-00565-z

Zhang, M., Cao, X., Jia, Q., & Ohlrogge, J. (2016). FUSCA3 activates triacylglycerol accumulation in Arabidopsis seedlings and tobacco BY2 cells. Plant Journal, 88(1), 95–107. 10.1111/tpj.13233

Zhao, Y., Chang, X., Qi, D., Dong, L., Wang, G., Fan, S., Jiang, L., Cheng, Q., Chen, X., Han, D., Xu, P., & Zhang, S. (2017). A novel soybean ERF transcription factor, gmERF113, increases resistance to *Phytophthora sojae* infection in soybean. Frontiers in Plant Science, 8, 299. 10.3389/fpls.2017.00299

Zhou, J. M., & Zhang, Y. (2020). Plant immunity: Danger perception and signaling. Cell, 181(5), 978–989). 10.1016/j.cell.2020.04.028

Zhou, Q., Galindo-Gonzalez, L., Manolii, V., Hwang, S. F., & Strelkov, S. E. (2020). Comparative transcriptome analysis of rutabaga (*Brassica napus*) cultivars indicates activation of salicylic acid and ethylene-mediated defenses in response to *plasmodiophora brassicae*. International Journal of Molecular Sciences, 21(21), 1–26. 10.3390/ijms21218381

Zhou, Q., Jayawardhane, K. N., Strelkov, S. E., Hwang, S. F., & Chen, G. (2022). Identification of Arabidopsis phospholipase A mutants with increased susceptibility to *Plasmodiophora brassicae*. Frontiers in Plant Science, 13. 799142. 10.3389/fpls.2022.799142

